# Host genetic control of succession in the switchgrass leaf fungal microbiome

**DOI:** 10.1101/2021.03.26.437207

**Authors:** A. VanWallendael, G. M. N. Benucci, P. B. da Costa, L. Fraser, A. Sreedasyam, F. Fritschi, T. E. Juenger, J. T. Lovell, G. Bonito, D. B. Lowry

## Abstract

Leaf fungal microbiomes can be fundamental drivers of host plant success, as they contain pathogens that devastate crop plants and taxa that enhance nutrient uptake, discourage herbivory, and antagonize pathogens. We measured leaf fungal diversity with amplicon sequencing across an entire growing season in a diversity panel of switchgrass (*Panicum virgatum*). We also sampled a replicated subset of genotypes across three additional sites to compare the importance of time, space, ecology, and genetics. We found a strong successional pattern in the microbiome shaped both by host genetics and environmental factors. Further, we used genome-wide association mapping and RNA-sequencing to show that three cysteine-rich receptor-like kinases were linked to a genetic locus associated with microbiome structure. These genes were more highly expressed in genotypes susceptible to fungal pathogens, which were central to microbial covariance networks, suggesting that host immune genes are a principal means of controlling the entire leaf microbiome.

## Main

Microbial communities perform essential functions for their host organisms in all branches of life. In some systems, hosts can tightly control the microbes with which they form symbioses^1,2^. In other systems the composition of the microbiome is more governed by ecological interactions such as the order of species arrival or abiotic conditions during colonization^3,4^. A key goal of microbial evolutionary ecology is to determine how both host and non-host factors influence microbiome assembly^5^, particularly in natural settings where host influence is more challenging to study.

Communities that colonize available niches in the process of succession follow certain predictable ecological patterns. Early-arriving species are typically those with effective long-range dispersal, while the climax community is dominated by species that can more effectively use resources under competition^6^. While these broad patterns are generalizable, the composition of any particular successional community depends greatly on both the habitat colonized and interspecific interactions such as priority effects, where the order of arrival of taxa governs the success of later arrivals^7^. While most successional theory is based on studies in macro-scale organisms, the principles of succession are evident in microbial communities as well, but on a more rapid time scale^8,9-10^.

In the case of microbiomes, host factors governing microbial succession must also be considered. Since the composition of the microbiome can greatly impact host fitness, it can be evolutionarily beneficial for the host to play a role in the successional process, encouraging mutualist colonization while dispelling pathogens as the community assembles. Hosts express genes that influence colonizing microbes through several means, including immunity, morphological adaptations^11^, and chemical exudation^12^. While the immune system is often effective at preventing detrimental infections, immune receptors may recognize and exclude beneficial microbes if elicitors are structurally similar to a pathogen, so specific immunity can have wider impacts on the microbiome^13^. Hosts require finely calibrated mechanisms for attracting beneficial microbes without attracting pathogens in a constant coevolutionary push and pull.

The phyllosphere microbiome, consisting of the microbes on and inside the plant leaf, comprises diverse taxa that impact plant health and productivity^14–17^. Leaf fungi in particular are common plant pathogens^18^, but non-pathogenic taxa may perform beneficial functions for the host, including nutrient uptake and pathogen antagonism^19–24^. Since the phyllosphere microbiome of perennial plants is reassembled at the start of each growing season in freshly-sprouted tissues,^25,26^ it may show similar patterns to macro-scale secondary successional communities.

We hypothesized that the phyllosphere fungal microbiome develops seasonally as a successional community controlled by environmental factors, host genetics, and interspecific fungal-fungal associations. We used amplicon sequencing to compare the relative importance of these factors in the phyllosphere fungi of a replicated diversity panel of switchgrass (*Panicum virgatum*^*27*^). We tested whether communities change directionally and whether the trajectory of succession differed across switchgrass genetic subpopulations and across different growing sites. Additionally, we sought to uncover whether specific genetic loci underlie host control of the microbiome through GWAS and RNA-sequencing analyses. Finally, we investigated the roles of specific fungal taxa in the microbiome through network analysis. Specifically, we aimed to determine whether known switchgrass leaf pathogens^28^ covary with nonpathogenic symbionts, or are peripheral to microbial communities.

## Results

### Succession varies across host subpopulations and planting sites

Switchgrass is a highly genetically diverse perennial grass native to North America, and both plant traits and switchgrass-microbe interactions vary across its range^27–29^. We leveraged this diversity to assess the difference in microbial communities across the three main switchgrass subpopulations by randomly selecting 106 genotypes from a diversity panel^27^ planted at our focal site, the Kellogg Biological Station (KBS), MI, USA. Of these, 28 genotypes were from the Midwestern subpopulation, 38 from the Atlantic, 31 from the Gulf, and 9 showed signs of admixture between groups (Intermediate). These subpopulations differ in morphological and ecological characteristics, so we expected that fungal succession would differ as well across subpopulations. We examined succession over time by sampling leaf tissue from each plant at five time-points, then sequencing the Internal Transcribed Spacer (ITS) region of the phyllosphere-associated fungi in and on the leaf. After quality filtering, we clustered 47.8 million ITS reads to 6756 fungal Operational Taxonomic Units (OTUs) that varied across genotypes and over time.

To determine the directionality of successional changes in the microbiome, we visualized community differences with nonmetric multidimensional scaling (NMDS). NMDS accurately preserved sample distances in reduced dimensions (Stress = 0.102; Figure S1), and revealed clear temporal community structure. NMDS1 clustered closely with the date of collection, while NMDS2 clustered more with host genetic subpopulation (Figure 1a). Notably, the first sampling date was highly distinct from the later time points, showing greater variation within that time point, as well as divergence from later time points (Figure 1). To explore the statistical significance of visual patterns of succession, we used PERMANOVA. In addition to sampling date (Day Of Year: DOY) and subpopulation, we compared the community differences on leaves that showed clear pathogen disease symptoms to uninfected leaves from the same plant. All terms we tested had significant effects on community structure, but differed greatly in their explanatory power (Table 1). At the focal site, KBS, collection date (DOY) explained the greatest amount of variation (19.4%), followed by genetic subpopulation (5.7%) and infection (1.2%), and there was a significant date-by-subpopulation interaction.

**Figure 1:**
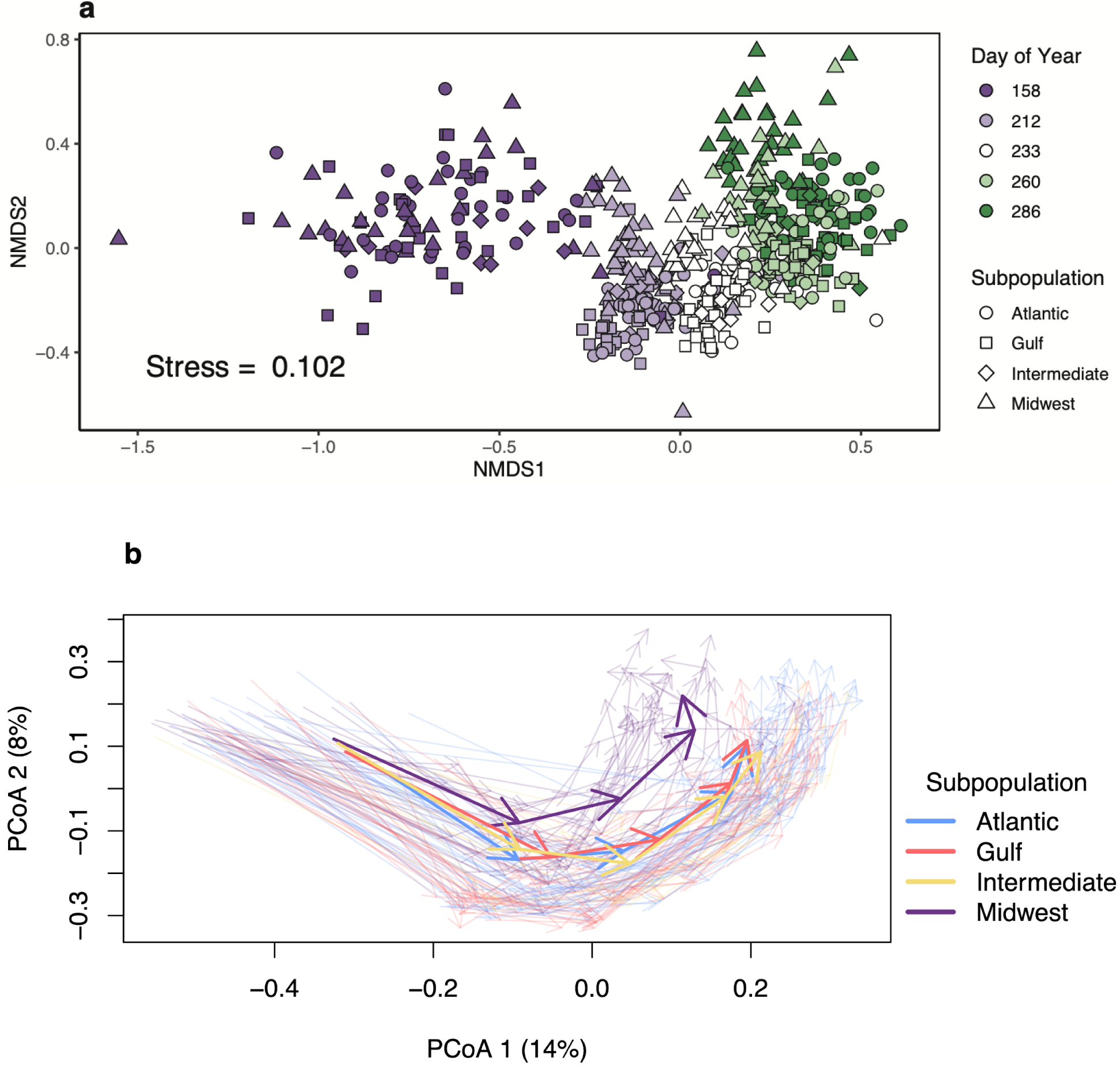
Structural and successional change in the leaf phyllosphere community shown by two methods. Each point represents an individual plant sampled from the experimental plot at Kellogg Biological Station, MI. **a**. NMDS (Non-metric Multidimensional Scaling). Dates are shown as Day of Year (DOY). Points are colored by DOY, and switchgrass subpopulations as shapes. NMDS does not estimate variation explained by individual axes, but uses nonparametric relationships to minimize a function of the difference between the representation and actual multidimensional distances called “stress.” **b**. Trajectory plots of principal coordinates of community distances. Transparent arrows represent individual switchgrass genotypes sampled over the five dates shown in **a**, and colors show genetic subpopulations. Solid colored arrows show mean subpopulation trajectories.

**Table 1:** PERMANOVA of a Bray-Curtis community dissimilarity matrix for genotypes at the focal site (n=106) and those replicated across sites (n=8). DOY (day of year) is sampling date, Subpopulation indicates genetic group, Infection indicates whether or not the sampled leaf had fungal disease symptoms, and Site indicates planting site. Terms are shown with sequential effects. All terms were significant with alpha < 0.05. At the focal site KBS, the model explained 28.2% of the total variation, whereas the multi-site test explained 43.2%.

| KBS Sequential |  |  |  |  |  | Multi-Site Sequential |  |  |  |  |  |
| --- | --- | --- | --- | --- | --- | --- | --- | --- | --- | --- | --- |
|  | Df | SumOfSqs | R2 | F | Pr(>F) |  | Df | SumOfSqs | R2 | F | Pr(>F) |
| **DOY** | **1** | **15.6** | **0.194** | **145.53** | **0.001** | **Site** | **3** | **4.6** | **0.296** | **10.95** | **0.001** |
| **Subpopulation** | **3** | **4.6** | **0.057** | **14.22** | **0.001** | **DOY** | **1** | **1.1** | **0.072** | **8.01** | **0.001** |
| **Infection** | **1** | **0.9** | **0.012** | **8.68** | **0.001** | **Subpopulation** | **2** | **0.7** | **0.046** | **2.54** | **0.001** |
| **DOY:Subpopulation** | **3** | **1.6** | **0.019** | **4.83** | **0.001** | **Infection** | **1** | **0.3** | **0.018** | **2.04** | **0.014** |
| Residual | 537 | 57.6 | 0.718 |  |  | Residual | 63 | 8.9 | 0.568 |  |  |
| Total | 545 | 80.3 | 1.000 |  |  | Total | 70 | 15.6 | 1.000 |  |  |

In order to directly test the differences in succession across subpopulations, we modeled changes in the multidimensional representation of fungal communities as directional trajectories^30^. Across the season, fungal communities on individual plants showed parallel changes over time, with almost no reversals to earlier community states (Figure 1b), strongly indicating a successional pattern. Switchgrass genetic subpopulations differed in both mean trajectory length (df = 3, F = 2.786; *p* = 0.0453) and mean overall direction (df = 3, F = 3.677; *p* = 0.0151). While little subpopulation difference is evident at the beginning of the season, climax fungal communities were markedly different in the Midwestern population, which showed the greatest divergence from others in trajectory direction (Figure 1b, Midwest-Atlantic; Tukey’s HSD = 0.013, *p* = 0.050). This provided initial evidence that, while fungal dispersal is similar across plant subpopulations, host plants influence the climax state of fungal communities.

Fungal microbiomes can be greatly influenced by environmental factors in addition to host factors. Therefore, we compared succession across environments by selecting a subset of eight plant genotypes replicated in three additional sites across a latitudinal gradient in the USA. From north to south, these field sites were Columbia, MO; Austin, TX; and Kingsville, TX (Figure S2). We sampled at each site at three time points, standardized by phenology to account for seasonal differences across sites (Figure 6). At most sites, collection date correlated with both NMDS1 and NMDS2 (Figure 2; stress = 0.103). However, the northern and southern sites were divided on a diagonal line orthogonal to collection date. The northern sites KBS and Columbia, MO formed one cluster, while the southern sites, Austin, TX and Kingsville, TX formed another (Figure 2). Differences across sites accounted for 29.6% of the variation in community dissimilarity across sites, but sampling date, subpopulation, and leaf infection also structured the community to a lesser extent (Table 1). While succession may show temporal patterns in southern sites, the composition of fungal communities on leaves is largely distinct.

**Figure 2:**
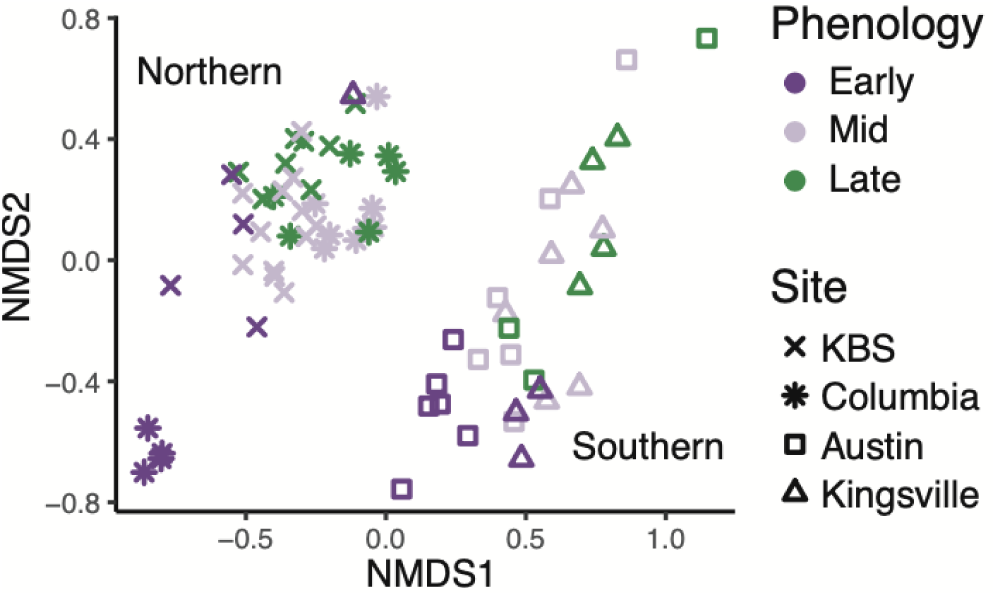
Site-specific changes in microbial communities shown by NMDS (Non-Metric Multidimensional Scaling). Genotypes were sampled at four sites. From north to south: KBS, MI; Columbia, MO; Austin, TX; and Kingsville, TX. Northern sites are shown by symbols, and southern by open shapes. Color indicates phenological stage sampled, “Early” samples were taken just after emergence, “Mid” samples were taken during seed development, and “Late” samples were taken after senescence began.

### Host genetic subpopulations support divergent fungal communities

Beyond differences at the level of subpopulations, we expected that within-subpopulation genetic differences would impact fungal diversity. To further examine genetic differences over time, we compared host genetic distances to fungal community differences between plants at the focal site, KBS. Genetic distances, calculated as Nei’s distance using 10.8 million single-nucleotide polymorphisms (SNPs)^31^, revealed that host population genetic structure largely matched the three major switchgrass genetic groups observed previously: ‘Gulf’, ‘Atlantic’, and ‘Midwest’^27^ (Figure 3a). These three subpopulations are deeply diverged and serve as discrete gene pools within which we tested for host-driven fungal community divergence. Fungal community distances, calculated as Bray-Curtis community dissimilarity, varied across sampling dates, but largely recapitulated the genetic structure of switchgrass (Figure 3bc). Notably, Mantel tests showed that fungal community structure was most closely correlated with host genetic structure at DOY 260, when most plants had set seed (r = 0.453), but declined as senescence progressed (Figure 3d). While we anticipated some degree of genetic influence, subpopulations were even more highly structured than expected, with almost half of the variation in fungal community distance explained by genetic distance when plants are setting seed (DOY 260).

**Figure 3:**
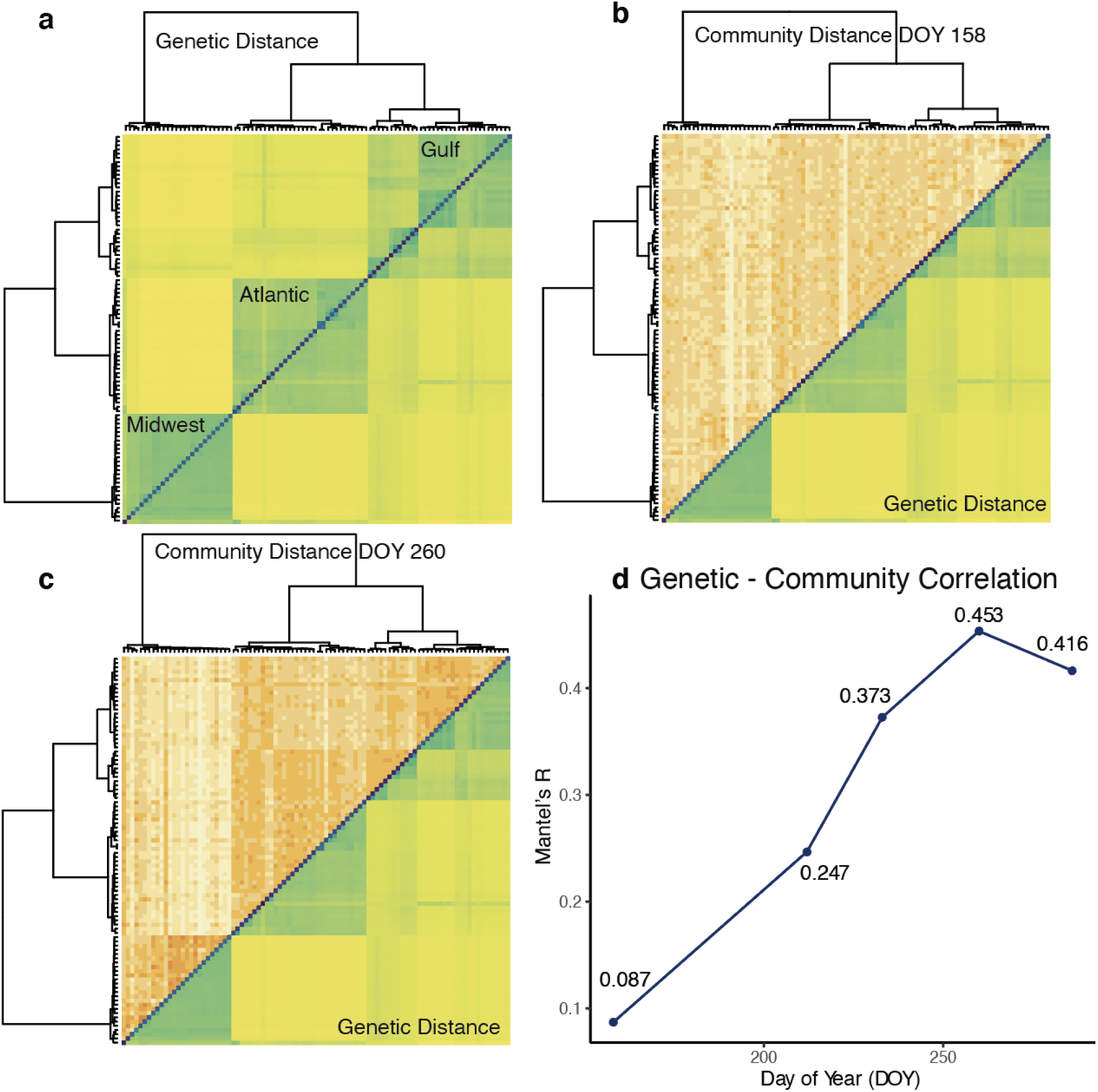
Genetic and fungal community pairwise distance matrices at Kellogg Biological Station, MI. **a**. Pairwise genetic distance (*π*) for all samples. Samples are ordered by hierarchical clustering. **b-c**. Pairwise community distance (Bray-Curtis) for all samples, shown in the same order as genetic distances, for two sampling times, DOY 158 and DOY 260. Other sampling times shown in Supp. **d**. Values of Mantel’s r shown indicate correlation between distance matrices for genetics and fungal communities at each time point. P < 0.01 for all tests.

Such tight host-microbiome genetic diversity associations imply a heritable genetic basis of fungal community dynamics by their plant hosts. To identify the genetic loci that might underlie this pattern, we calculated genome-wide associations (GWA) for microbiome structure. We used the second NMDS axis at DOY 260 from the above analysis (Figure 1; Figure S3) to represent microbiome structure, since it showed the greatest clustering with subpopulation (Figure 1a, 3c) and controlled for large-scale host genetic structure by including a single variate decomposition of pairwise genetic distance as a covariate in the linear models. We found several loci associated with the phenotype at a 5% false discovery rate (FDR), but the GWA showed an excess of low p-values (quantile-quantile plot: Figure S4) so we used a more conservative Bonferroni-corrected threshold to identify significant SNPs (Figure 4a). This threshold revealed only one SNP on chromosome 2N significantly associated with microbiome structure, Chr02N_57831909. This SNP is closely linked to several genes in the switchgrass v5.1 genome annotation (Figure 4bc). The three closest genes are homologous to receptor-like kinases (RLKs) annotated in the closely related *Panicum hallii* (two copies of *cysteine-rich receptor-like protein kinase 6; XP_025800480*.*1 & XP_025800481*.*1*, and one copy of *cysteine-rich receptor-like protein kinase 10; XP_025801715*.*1)*. This class of RLKs is diverse in plants, but is known to contain many immune receptors^32^, indicating a potential role for these genes in host control of fungi.

**Figure 4:**
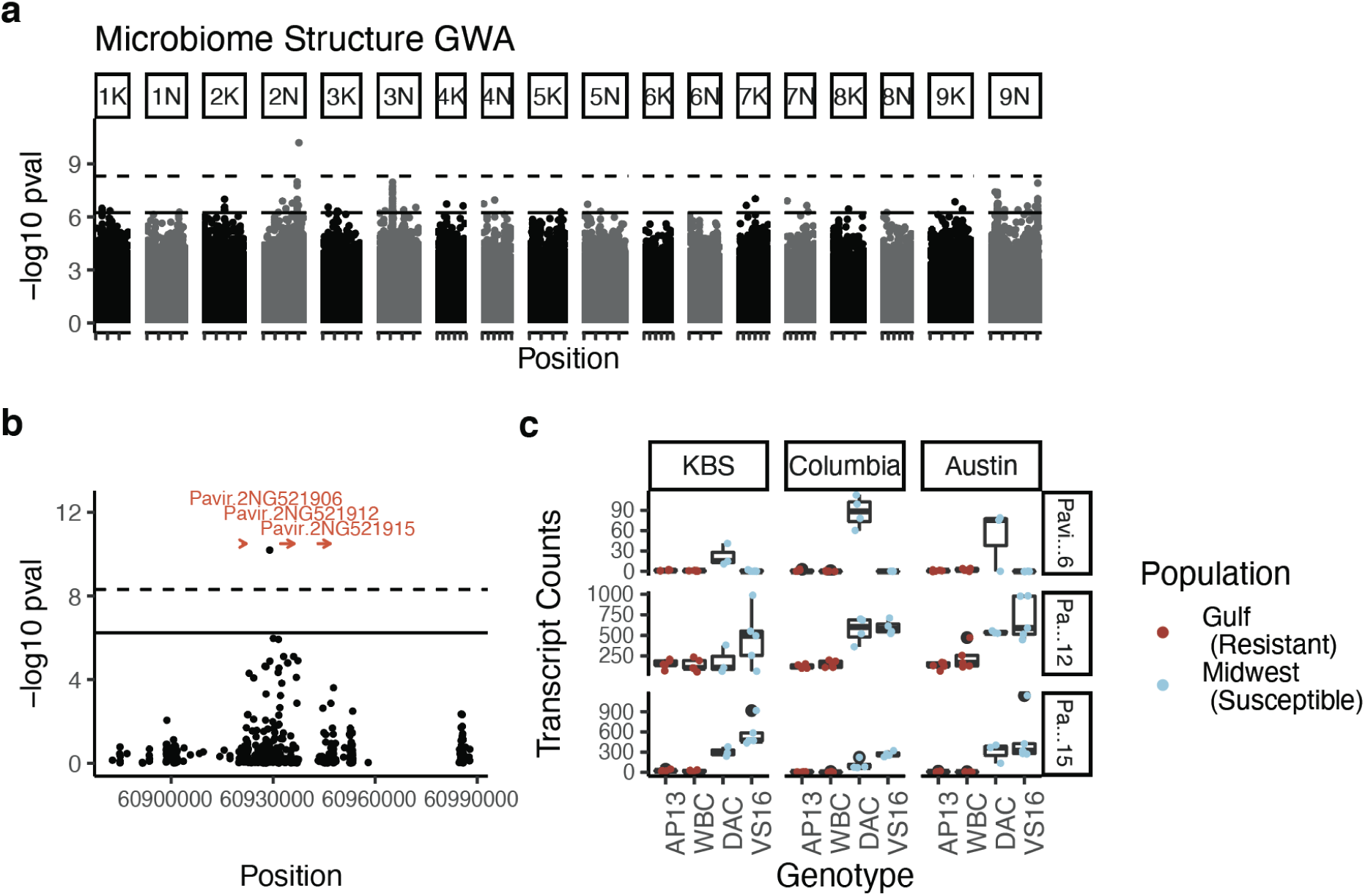
**a**. Genome-wide associations of microbiome structure (NMDS2 at DOY 260). The lower solid line shows the 5% False Discovery Rate threshold, and the upper dashed line shows the Bonferroni-adjusted alpha threshold for SNPs associated with the microbiome. **b**. Outlier region on Chromosome 2N with nearby genes shown in red. **c**. Expression-level differences for genes shown in **b**. Leaf tissue for these samples was collected as part of a different study, performed at three of the same sites we used. Transcript counts are scaled differently in each gene facet.

We corroborated the importance of these candidate genes by comparing their expression levels in divergent genotypes at the three of the four sites where phyllosphere experiments were conducted, KBS, Columbia, and Austin. In each site, we sequenced leaf tissue RNA from multiple biological replicates (n >= 3) from four genotypes: two that are susceptible to leaf fungal pathogens (Midwest upland VS16 & DAC) and two which were generally resistant (Gulf lowland WBC & AP13)^28^. Consistent with host-gene driven variation in fungal community assembly, all three candidate genes were much more highly expressed in susceptible than resistant genotypes (Wald tests; Table 2). These differential genotype-specific patterns of expression were very similar across planting sites (likelihood ratio test for ecotype ✕ site interaction, p = 0.354). The pattern of these receptors being more highly expressed in pathogen-susceptible plants may seem counterintuitive, since many RLKs are immune receptors. However, this can often occur when pathogens produce effector proteins that target immune receptors^33^. Necrotrophic fungi in particular can benefit by over-inducing plant immune receptors to initiate programmed cell death,^34,35^ then feeding on dead plant tissue.

**Table 2:** Wald test for expression differences in three candidate genes between susceptible (upland) and resistant (lowland) ecotypes.

|  | Base Mean | log Fold Change | SE | W | p-value |
| --- | --- | --- | --- | --- | --- |
| Pavir.2NG521906 | 12.91 | 3.22 | 0.81 | 3.95 | 7.89e-05 |
| Pavir.2NG521912 | 313.63 | 1.99 | 0.16 | 12.25 | 0.00e+00 |
| Pavir.2NG521915 | 189.09 | 5.33 | 0.30 | 17.94 | 0.00e+00 |

### Yeasts, pathogens, and mycoparasites are core phyllosphere microbiome members

Given the large differences in leaf pathogen susceptibility across switchgrass subpopulations, we sought to determine the influence of pathogenic fungi on other members of the fungal microbiome. We examined the taxonomic relationships of the 7392 OTUs in our dataset through a hybrid method that compares matches across fungal databases and BLAST (Basic Local Alignment Search Tool) hits^36^. We identified 6756 OTUs as fungi, 633 as plant, and 3 OTUs as metazoan. We performed NMDS and PERMANOVA analyses using the full community, but focused our taxon-specific analyses on OTUs at the focal site that were present at high occupancy across time and showed relatively high abundance, often referred to as the “core” microbiome^37^. This group consisted of 128 OTUs, the majority of which were Dothideomycetes (43.5%) and Tremellomycetes (28.7%; Table S1). We assigned each of the core OTUs to a functional guild when possible using published literature (Table S1). Of the core group, 23 OTUs were grass pathogens and 9 were documented pathogens of other plants. Four were known mycoparasites, fungi that prey upon other fungi. 3 were generalist decomposers or had an unclear functional guild, and the remaining 52 were yeasts or yeast-like fungi. Compared to fungal species in soil, these taxa were especially enriched for grass pathogens and yeasts, and contained much fewer sabrobes^38^.

To investigate how these functional guilds associate, we built covariance networks using OTU relative abundances at each time point (Figure 5a). We summarized network statistics across functional guilds to show that known grass pathogens are central to covariance networks, with high betweenness centrality (extent to which a node lies on paths connecting other nodes) and degree (overall number of connections; Figure 5e). Standard deviation was high within this group, however, reflecting seasonal and within-group differences. Yeasts, in contrast, showed higher modularity (compartmentalization; Figure 5e). This indicates that, while yeasts are overall more speciose in the core microbiome, they covary less with the rest of the microbial community than pathogens. Since yeasts are thought to be mostly commensal inhabitants of the outer leaf surface^39^, this difference may reflect their ecological or spatial niche.

**Figure 5:**
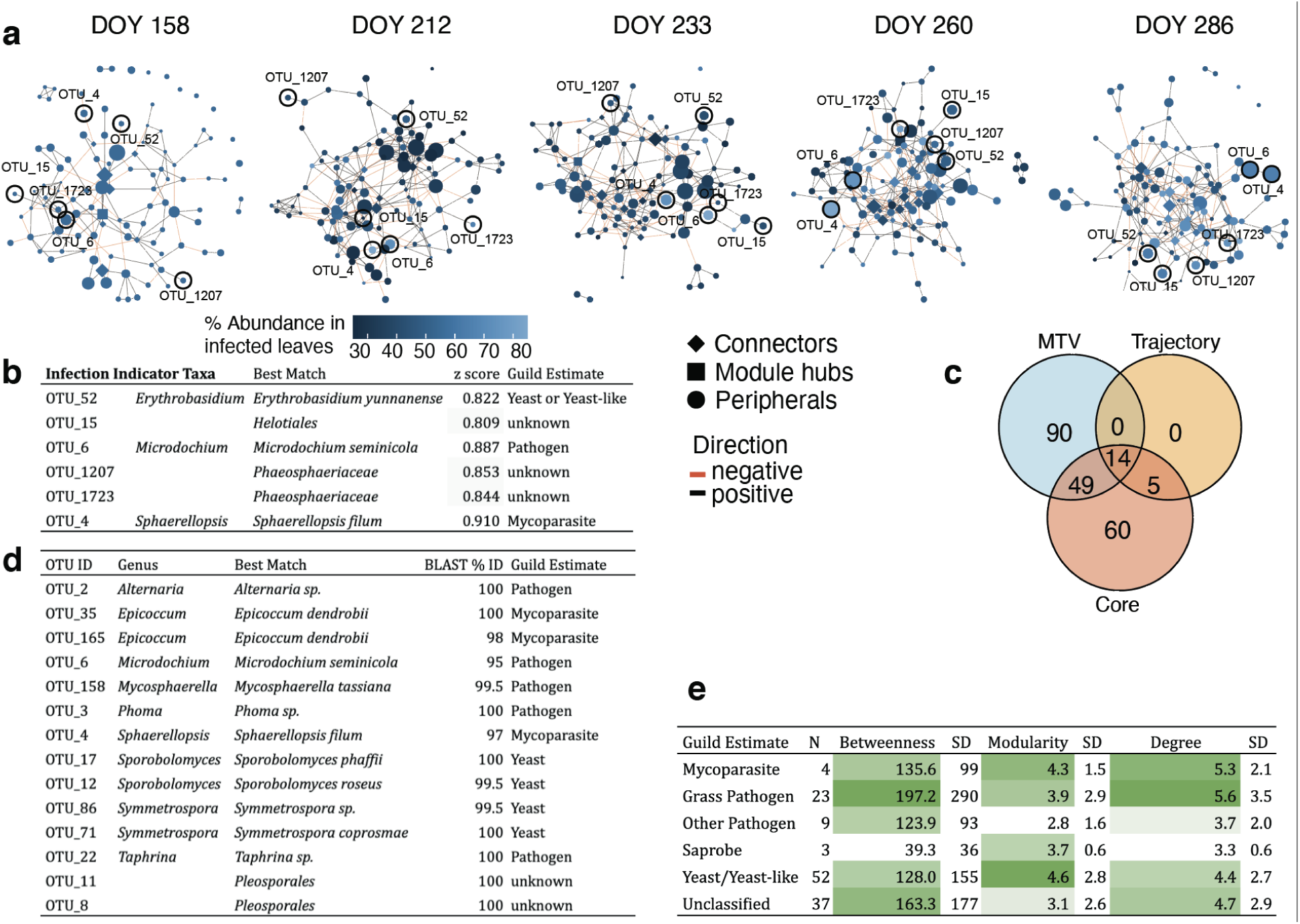
**a**. Covariance networks of core OTUs over time. Nodes are colored by each OTU’s relative abundance in infected leaves with visible symptoms. The shape of the node denotes network position, defined by Zi-Pi ratio. Edges are colored by the covariance sign. **b**. Infection indicator taxa, including best taxonomic match and z-score for indicator analysis. **c**. Number of OTUs identified as important by several methods: MTV-LMM analyses that indicate time-dependent OTUs, OTUs that impact the successional trajectory, and core OTUs with high occupancy-abundance. **d**. Taxonomic information for the fourteen OTUs identified in all three analyses in **c**. Best match denotes the lowest taxonomic level confidently identified for each OTU using BLAST. Guilds were estimated based on published studies, references are in Table S1. **e**. Network statistics for fungal guilds, calculated as mean values across all time points, with standard deviation (SD).

In addition to varying among functional groups, OTU covariance also changed over time (Figure 5a, S6). To identify positive or negative covariance temporal patterns within network members, we generated a Class-level heatmap showing the proportion of edges linking OTUs within or between each Class at each time point (Figure S5). Due to the high proportions of Dothideomycetes (mixed guilds) and Tremellomycetes (yeast) in the core, the majority of edges at every time point were within (38.3-48.6%) and between (9.9-14.5%) OTUs in these classes. While the proportion of positive edges maintained more or less stable with time between OTUs in the Dothideomycetes (from 20.0 to 23.9%) and Tremellomycetes (from 23.0 to 17.4%) or within the two classes (from 3.6 to 2.8%), negative edges between classes increased from DOY 158 (6.7%) to DOY 233 (9.7%) and DOY 286 (11.7%). This may indicate competition for host resources between these two classes of fungi, resulting in more spatially heterogenous distributions in the late season.

While patterns in this core group reflected major changes in fungal microbiome, we used several alternate methods to identify important OTUs. In addition to the core, we modified trajectory analyses (Figure 1b) by computationally removing each OTU from the analysis and calculating the change in the overall community trajectory^30^. Nineteen OTUs significantly impacted trajectories when they were removed, all of which overlapped with the core group (Figure 5c). To examine priority effects, we used microbial temporal variability mixed linear models (MTV-LMMs), which identify taxa for which variation in earlier time points explains variation in later points^40^. Of the 153 OTUs we found in this analysis, 49 overlapped with the core, and 14 with both the core and trajectory analysis (Figure 5c). The fourteen OTUs that were identified as important using all three methods (Figure 5cd) included taxa from several putative functional guilds, including yeasts, pathogens, and mycoparasites. Network connections confirmed mycoparasitic interactions; we found a negative relationship between putative plant pathogens *Mycosphaerella tassiana* and *Microdochium seminicola* and mycoparasitic *Epicoccum dendrobii*. In addition, we used indicator species analysis to identify OTUs that were overrepresented in leaves with fungal disease symptoms (Figure 5a). Although the major fungal pathogen of switchgrass, *Puccinia novopanici*, was not identified as a core taxon, indicator species analysis showed that putative mycoparasite *Sphaerellopsis filum* is present in the core, and significantly associated with fungal infection symptoms (Figure OTU_4; Figure 5a).

Our analyses mostly identified the taxa that were abundant across samples. Rare taxa can be important in microbial community functioning ^41^, but their role in overall ecological patterns is less clear and more challenging to study. Therefore, we only examined rare taxa that we expected *a-priori* to play an important ecological role. *Claviceps* species were present in 119/760 samples, and were more highly abundant in the early season. *Claviceps* species produce alkaloid compounds that deter grazing^42^, so this endophyte may play a role in protecting young grass shoots. *Metarhizium*, a related genus, was present at low abundances in 43/760 samples in the Columbia, MO and KBS, MI sites. *Metarhizium* species are insect-pathogenic fungi^43^, so may provide a similar protective role.

## Discussion

Our results show strong support for the importance of time, geographic location, and host genetics in influencing the switchgrass phyllosphere microbial succession over the growing season. We found evidence for clear successional dynamics that were consistent in direction across growing sites, but were distinct in community composition. Fungal communities were different across host genetic subpopulations, a pattern that may be driven in part by variation at three linked immune receptors. While there is great taxonomic diversity in leaf fungal communities, a few highly abundant taxa, many of which are pathogens, play a disproportionate role in shaping community progression and likely influence plant host traits.

### Succession varies across host subpopulations and planting sites

Viewing the switchgrass leaf microbial community through the lens of succession allowed us to delineate ecological patterns in these communities. Multidimensional scaling representations of the leaf communities at the focal site revealed a clear clustering by date of collection on the first NMDS axis (Figure 1). This indicates that, as we predicted, date of collection is an important source of variation in the switchgrass leaf fungal community. Further, measuring the trajectories of these communities showed that succession is both directional and deterministic, since no samples showed negative trajectories (reversals of succession) by the end of the season, and most samples followed a similar trajectory (Figure 1b). While the overall shape of trajectories was similar, the Midwestern population deviated from others, particularly in the late season. The Midwestern population is notable since we have previously shown that it is more susceptible to several fungal pathogens such as leaf rust (*Puccinia novopanici*) and leaf spot (*Bipolaris spp*.; ^28^; also see ^44^), and has on average an earlier phenology than the other population groups.^29^ Leaf microbiome relationships in this population are consistently distinct in this population, and may be linked to other traits such as cold-tolerance that also differ^27,29^.

In addition to temporal differences across subpopulations, the composition of fungal leaf communities differed markedly across geographic locations. This may be partially due to seasonality differences across the region we examined. The Kingsville, TX site did not experience freezing temperatures between 1989 and 2020 (NOAA weather service), so perennial grasses in the region may have living aboveground tissue year-round. Growing season length has been shown as an important factor in governing the abundance and diversity of endophytic fungi^45^, so it is unsurprising that we saw large differences across this latitudinal gradient. However, many other factors that influence fungal communities also differ across these sites, including precipitation regime, soil type, and surrounding vegetation, so further work is needed to determine if the growing season is truly the causal factor. Climate plays an important role in regulating plant endophyte diversity ^46^

### Host genetics influence community structure

We predicted that fungal communities would be impacted by host genetics as well as location. We found several lines of evidence for genetic control of the leaf microbiome. In addition to examining differing successional trajectories across subpopulations, we tested the covariance of genetic distance and fungal community differences using Mantel correlations. Genetic-fungal community correlations increased until DOY 260, then declined as host senescence began. Mantel tests are inappropriate for some ecological tests and often underestimate p-values, but can be useful for exploratory analysis of distance matrices^47^. When selecting samples for this study, we randomly chose equal numbers of samples from the two major switchgrass morphological ecotypes, upland and lowland switchgrass^48^ (Figure S2). Lowland switchgrass, which is more highly represented in Gulf and Atlantic subpopulations, is more resistant to several leaf fungal pathogens^28^, so subpopulation differences may be at least partially driven by differences in immunity across these genotypes. Since pathogens such as *Microdochium* and *Alternaria* were among the most abundant taxa in our samples, their differences across subpopulations may have driven overall community differences. In addition to immunity, however, subpopulations differ in other traits that may contribute to fungal colonization differences, such as leaf wax content ^49^, exudate concentration^50^, and phenology^29,48^, so microbiome differences may be responding to multiple host plant traits.

### A replicated receptor-like kinase is associated with fungal differences

We found one outlier SNP associated with microbiome structure. While there were several peaks in the Manhattan plot (Figure 5a), our analysis showed a strongly skewed distribution of observed versus expected p-values (Figure S4), indicating a risk of Type I errors. This is probably attributable to the low sample size in this GWAS. The influence of the identified locus is fairly strong, contributing to a clear decrease on NMDS axis 2 when the minor allele is present (MAF = 0.083; Figure S6). This SNP is not in Hardy-Weinberg equilibrium in switchgrass; we found only one minor-allele homozygote among our samples. This abnormal pattern may be attributable to structural variation at this locus. Switchgrass subpopulations vary widely in genome structure, which may result in alignment mismatches that resemble SNPs, particularly in regions with multiple gene copies^51^. Indeed, this region shows an elevated number of insertions and deletions compared to nearby sections of the 2N chromosome (Figure S7, data from ^27^). Given the strong association for this locus as well as the RNA-sequencing results, however, we expect that there is a true phenotypic association with the locus, but that it may be with a structural variant rather than a true SNP.

The three nearby genes we identified were replicated variants of a cysteine-rich RLK whose function has not been experimentally verified in *Panicum*. RLKs are one of the largest plant gene families, including over 600 members in *Arabidopsis*^*32*^. The best-studied of these is FLS2, which detects the bacterial flagellin protein and initiates an immune response cascade^52^. The three RLKs we identified show high sequence similarity to immune-related cysteine-rich RLKs in *Arabidopsis* and *Oryza*, and contain the “stress-antifungal domain” PF01657, which has been linked to salt stress as well as fungal responses when present in several proteins^53,54^. *Arabidopsis CRK5*, for instance, alters defense responses either through resistance to infection or programmed cell death, depending on how the gene is expressed ^55^. Similarly, the *Oryza* gene *LIL1* (Os07g0488400) improves fungal rice blast resistance when overexpressed ^56^.

### Pathogens & hyperparasites are important in succession

We used several methods to identify important taxa in the phyllosphere community. We used “core” microbiome analysis to identify OTUs that show high occupancy (presence across multiple samples within a time point^15^). We found that core taxa overlapped well with important taxa identified by MTV-LMMs and trajectory analysis. We can therefore be confident that this group of taxa is influential in the switchgrass phyllosphere (Fig. 5). Within this group, we identified several as pathogens, including *Alternaria, Mycosphaerella, Microdochium*, and *Taphrina*. It is challenging to assign functional guilds to symbiotic fungi, since their benefit or detriment to the host may depend strongly on phenology, abiotic conditions, and ecological interactions^57^. For instance, many endophytic fungi are commensal for most of the season, then shift to breaking down plant tissue as the host begins senescence^58^. Others may be weakly pathogenic, but may improve overall host fitness by enhancing nutrient uptake or preventing infection by more effective pathogens^22,59^.

Yeasts and yeast-like fungi were also well-represented in phyllosphere samples. Yeasts were historically thought to be dominant in the phyllosphere^60^, but this may have been an artifact of methods used. Yeasts are more easily culturable than filamentous fungi, and are therefore overrepresented in studies using cultures to measure fungal diversity. The exact relationship between yeasts and plant hosts is not totally clear, but they are typically thought to be mostly commensal symbionts, feeding on small amounts of sugars on the leaf surface^61^.

At the focal site, Tremellomycete yeasts and Dothideomycetes dominated the core microbiome and covaried negatively through time. This may be explained by different spatial distributions across samples; Tremellomyctes dominate some samples and Dothideomycetes other samples, but they rarely coexist. Priority effects, wherein early-arriving taxa gain advantage over late-arriving taxa, may therefore play a role in governing colonization in these taxa. Certain Tremellomycete yeasts have been shown to be potential biocontrol agents against pathogens, e.g. *Papiliotrema spp*.^*62*^, and others have been shown to be “hub” taxa or negatively connected with leaf pathogens, e.g. *Dioszegia spp*.^*63*^, both genera with high abundance in our focal site dataset.

One unexpected finding of our taxon-specific analysis was that two mycoparasites were identified as important taxa, *Epicoccum* and *Sphaerellopsis. Epicoccum* is an ascomycete genus comprising several species with noted antifungal properties^64,65^. The species we identified in this study, *Epicoccum dendrobii*, is being investigated as a biocontrol agent of the pathogenic anthracnose fungus *Colletotrichum gloeosporoides*^*66*^. Similarly, *Sphaerellopsis filum* has been observed infecting multiple species of *Puccinia* rusts^67,68^, and has been shown specifically to reduce switchgrass rust infection^69^. Another surprising finding was that switchgrass rust was not a core species, despite the fact that its disease symptoms are nearly omnipresent each year in the sites we studied^28^. Fungi in the Pucciniaceae family have an ITS sequence that differs substantially from general fungal primers used in this study, which we suspect resulted in reduced amplification of *Puccinia* rusts. However, the ubiquity of the *Sphaerellopsis* hyperparasite is an indication that *Puccinia* may be more prevalent than our sequencing data show, a speculation that is supported by the fact that *Sphaerellopsis* was identified by indicator species analysis as clearly overrepresented in leaves with rust infection.

## Conclusion

Switchgrass leaf fungal communities are highly diverse, and are influenced by both host and environmental factors. Succession occurs each season as communities are assembled through stochastic, environmental, and host-determined processes. Pathogenic fungi play a critical role in the switchgrass leaf phyllosphere community, determining both the trajectory of microbial community development and acting as central nodes in community networks. Host immune genes such as receptor-like kinases control pathogens directly, and the prevalent mycoparasites that prey on them indirectly. The plant genes that control pathogens may therefore provide a principal means by which plants influence changes in their fungal microbiome.

## Material and Methods

### Plant material

We collected switchgrass leaves from a diversity panel established for a separate study^27^. In brief, researchers planted arrays of 732 genotypes of switchgrass clonally replicated at over fifteen sites in the US and Mexico. These genotypes were collected from across the United States, grown in controlled conditions, then clonally split before replanting at all sites. Since 2018, they have been growing in 1.3 m spaced grids with minimal interference for weed control^27^. Researchers used Illumina HiSeq X10 and Illumina NovoSeq6000 paired-end sequencing (2×150bp) at HudsonAlpha Institute for Biotechnology (Huntsville, AL) and the Joint Genome Institute (Walnut Creek, CA) to sequence the genome of each individual. Sequence information for these samples is available on the NCBI SRA: Bioproject PRJNA622568. Lovell et al.^27^ called 33.8 million SNPs with minor allele frequency greater than 0.5%, 10.8 million of which we used in this study for genetic mapping.

We used two sampling strategies to assess temporal and geographic variation (Figure 6; Table S2). For temporal variation, we sampled leaf tissue from 104 genotypes from a diversity panel of switchgrass grown at the Kellogg Biological Station, MI field site at five time points during the 2019 growing season. To assess geographic variation, we collected eight randomly chosen genotypes representative of switchgrass genetic populations that were replicated in four sites that span the geographic range of temperate switchgrass populations: Kellogg Biological Station, MI; Columbia, MO; Austin, TX; and Kingsville, TX. At each site, we sampled the same eight genotypes at three time points (n = 96; Figure 1). Given that climate varies greatly over this latitudinal range, we standardized collection by phenology rather than date, focusing on switchgrass emergence, flowering, and senescence. Switchgrass genetic variation segregates into three main subpopulations that differ greatly in morphology and phenology^27^, so we compared fungal community responses over these populations.

**Figure 6:**
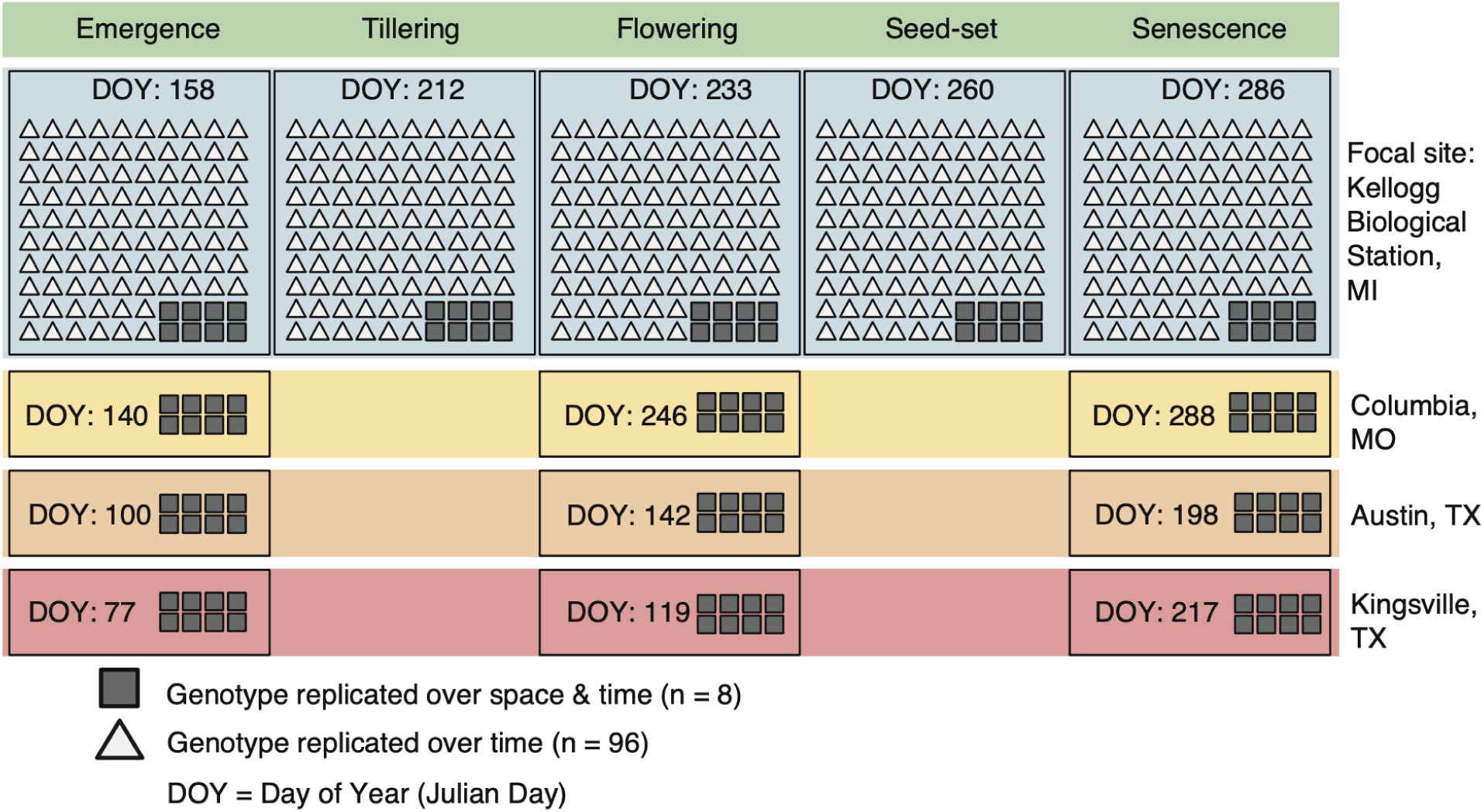
Sampling scheme. Each icon represents one sample taken. We sampled 106 genotypes at five time points at the focal site in Hickory Corners, MI, and 8 genotypes at the three other sites. We sampled roughly equal numbers of each subpopulation throughout. Sites are shown from northern (Kellogg Biological Station, MI) to southern (Kingsville, TX).

We used two sampling strategies to assess temporal and geographic variation. For temporal variation, we sampled 104 genotypes from the Kellogg Biological Station, MI field site at five time points, corresponding to the phenological stages of emergence, tillering, flowering, seed-set, and senescence (n = 530; Figure 6). To assess geographic variation, we selected a latitudinal transect of four of the fifteen sites to sample: Kellogg Biological Station, MI (42.419, −85.371); Columbia, MO (38.896, -92.217); Austin, TX (30.383, -97.729); and Kingsville, TX (27.549, -97.881). At each site, we sampled the same eight genotypes at three time points (n = 96; Figure 6; Figure S4). Since climate varies greatly over this latitudinal range, we standardized collection by phenology rather than date, focusing on switchgrass emergence, flowering, and senescence. At all sites we collected roughly equal numbers of genetic subpopulations.

For each plant at each time point, we collected 3 leaves. We haphazardly sampled leaves from the middle of the canopy; that is, leaves that were neither close to the base nor the flag leaf. To minimize external contamination, we sterilized gloves between plants, and collected directly into sterile 50mL tubes (UHP tubes, Fisher Scientific, Waltham, MA). Since we expected that the fungal community would be impacted by the dominant fungal pathogen, leaf rust, we collected 3 leaves with visible rust symptoms as well as 3 uninfected from the same plant when possible (n = 106), all of which were used in downstream analyses. We stored tubes on dry ice in the field and while being shipped, then at −80°C until extraction. For each day of sampling, we also collected a negative control, one tube opened to the ambient air for at least 10s. Samples were shipped overnight on dry ice to Michigan State University (MSU) for processing.

### ITS sequencing

We targeted the endophytic (inside the leaf) and epiphytic (on the leaf surface) fungi. To prepare leaves for DNA extraction, we used 4mm biopsy punches (Integra, Princeton, NJ) to produce ~21 leaf discs pooled across the three collected leaves. We sterilized biopsy punches between samples by soaking overnight in DNAaway (Thermo Fisher Scientific, Waltham, MA), then washed in DI water. We punched across the leaf blade to fully represent the spatial diversity in leaves. We homogenized leaf tissue by grinding with two sterile 3.175 mm stainless steel ball bearings. We placed sealed sterile 1.5 ml tubes with bearings and leaf discs into liquid nitrogen for 10s, then homogenized in a Mini-G bead beater (SPEX sample prep, Metuchen, NJ) for 60 s at 1500 rpm.

To extract DNA, we used Qiagen Plant Maxi kits, following the manufacturer’s instructions (Qiagen, Hilden, Germany). This method yields large amounts of plant DNA in addition to fungal, so we used primers for the fungal ITS (Internal Transcribed Spacer) rDNA region. We performed library preparation for ITS using the ITS1f (5’-CTTGGTCATTTAGAGGAAGTAA-3’) and ITS4 (5’-TCCTCCGCTTATTGATATGC-3’) primers. We used a three-step amplification process to amplify the target region, add adaptors, and add barcodes for multiplexing as previously reported by Benucci et al.^70,71^ PCR amplification steps and reagents are included in the supplement (Table S3). We normalized DNA concentrations using SequalPrep normalization kits (Thermo Fisher Scientific, Waltham, MA), concentrated libraries using Amicon Ultra 0.5 mL 50K centrifugal filters (EMD Millipore, Burlington, MA), and removed primer-dimers with Ampure magnetic beads (Beckman Coulter, Brea, CA). We randomized samples across plates, then pooled them into 3 libraries for sequencing. We used four levels of negative controls to check for contamination at different steps: field controls that were exposed to air at each sampling point, DNA extraction controls, library preparation controls, and a synthetic mock community^72^, resulting in a total of 672 samples that included 59 controls.

We sequenced DNA using Illumina MiSeq 300bp paired-end v3 600 cycles kit in the MSU genomics core facility. Sequencing yielded 84.7 M total reads, and high quality data across samples. Across three multiplexed libraries, 74.9% of reads had quality scores above 30 (Phred), with an average of 110 K reads per sample (ranging from 110 reads in negative controls to 199 K reads in samples). After quality filtering, 47.8 M reads remained. We used a 97% clustering threshold for identifying OTUs (Operational Taxonomic Units), resulting in 7963 OTUs across 761 samples.

### RNA Sequencing

Vegetatively propagated plants from four genotypes were grown in three sites (KBS, MI; Austin, TX; and Columbia, MO). Two genotypes, AP13 and WBC, fit in the Gulf population group, and are generally resistant to leaf fungal pathogens^28,44^. The other two are more closely related to the Midwest population, and are more susceptible to leaf pathogens^28,44^. Leaf tissue was harvested and immediately flash frozen in liquid nitrogen and stored at −80°C until further processing was done. Each harvest involved at least three independent biological replicates. High quality RNA was extracted using standard Trizol-reagent based extraction^73^. RNA-Seq libraries were prepared using Illumina’s TruSeq Stranded mRNA HT sample prep kit utilizing poly-A selection of mRNA. Sequencing was performed on the Illumina HiSeq 2500 sequencer using HiSeq TruSeq SBS sequencing kit. Paired-end RNA-Seq 150-bp reads were quality trimmed (Q ≥ 25) and reads shorter than 50 bp after trimming were discarded. High-quality sequences (404.4 M reads) were aligned to P. virgatum v5.1 reference genome using GSNAP v.2019-06-10^74^ and counts of reads uniquely mapped to annotated genes (371.8 M reads) were obtained using HTSeq v.0.11.2^75^.

### Bioinformatics

We analyzed ITS sequences on the MSU HPCC (High-Performance Computing Center) with *qiime* v1.9.1^76^, *fastqc* v0.11.7^77^, *cutadapt* v2.9^78^, *CONSTAX2*^*79*^and *usearch* v11.0.667 ^80^. We demultiplexed sequencing reads using *split_libraries_fastq*.*py* in *qiime1*, then checked for sequencing errors with *fastqc*. We removed barcodes with cutadapt, and filtered fastqs with *USEARCH using the* -fastq_filter option with arguments: 1 expected error (-fastq_maxee 1.0), truncation length of 200 (-fastq_trunclen 200), and no unidentified bases (-fastq_maxns 0)^81^. We clustered 97% OTUs with the UPARSE algorithm^82^ through the -cluster_otus option, with singletons discarded (-minsize 2). We assigned taxonomy to OTUs using CONSTAX2^36^, which improves OTU identifications using a consensus algorithm between RDP^83^, SINTAX^84^, and BLAST classifications^36^.

### Statistical Analyses

We performed downstream analyses in R v4.0.3 ^85^ using the packages *decontam*^*86*^, *vegan*^*87*^, *phyloseq*^*88*^, *vegclust*^*89*^, and *metagenomeseq*^*90*^. We used *decontam* to remove contaminants by pruning OTUs that were overrepresented in negative controls, then normalized read depth with functions in the *metagenomeseq* package. Of 7963 OTUs we clustered, 162 were identified as contaminants and removed from analyses (identifiable contaminants removed are shown in table S4). All contaminants showed low abundance and were evenly spread across negative controls, indicating that fungal contamination was minimal in this study.

### Successional dynamics

We visualized community structure using nonmetric multidimensional scaling (NMDS), which represents the multivariate structure of a community in reduced dimensions (Shepard plot in Supplemental Figure 1). We first used a Hellinger transformation to standardize across samples using the *decostand* function, then performed NMDS with *metaMDS*, both in the *vegan* package. We also used permutational analysis of variance to assay the relative importance of various factors in structuring the fungal community implemented through the *adonis2* function in *vegan*.

To test the importance of historical contingency in temporal community changes, we used a Microbial Temporal Variability Linear Mixed Model (MTV-LMM^40^). The MTV-LMM assumes that temporal changes are a time-homogenous high-order Markov process, and fits a sequential linear mixed model to predict the abundance of taxa at particular time points ^40^. For each taxon, we calculated “time explainability”, a metric of the degree to which variation in later time points is explained by variation in earlier points^40^. We fit linear mixed models for each OTU present across multiple time points, and used a Bonferroni-corrected alpha to identify taxa that exhibit significant temporal contingency.

In addition, we examined individual and population-level succession using trajectory analysis^30^. Trajectory analysis transforms multivariate community changes to two-dimensional trajectories, for which parameters of individual community changes can be compared. We calculated mean trajectories for communities in each subpopulation, then used ANOVA to test for trajectory differences across subpopulations. We then used a permutational method to discover OTUs that substantially impact succession. We computationally removed each OTU from our dataset, recalculated mean population trajectories, then compared to the original trajectories. We then used the *trajectoryDistances* function in the *vegclust* package to calculate the degree to which removing each OTU altered the overall community trajectory^89^.

Conceptualization of ecological communities as trajectories has a long history in ecology^91^, but explicit modelling of trajectory parameters has been challenging until relatively recently^30,92,93^. This approach utilizes statistical methods that are typically applied to movement in geometric space^94^ to compare movement by a community in multidimensional space^30^. While trajectory analysis has not been applied to changes in microbial communities to our knowledge, other researchers have used the method to understand succession in Amazon forest communities after land-use change^95^, and Iberian forests after fires^96^.

### Genetic associations

To specifically measure the overall microbiome variation explained by genetic structure, we examined the covariation of genetic distance and fungal community distance using Mantel tests. We calculated genetic distance as the number of pairwise SNP differences between each sample (Nei’s distance, π). We used the switchgrass GWAS SNP dataset^31^, which features 10.8 million high-confidence SNPs with minor allele frequency (MAF) > 0.05, and calculated distance with the *dist*.*genpop* function in *adegenet*^*97*^. For microbiome community differences, we used Hellinger-transformed Bray-Curtis distances calculated with the *decostand* function in *vegan*. We performed Mantel tests with 999 permutations using the *mantel* function in *vegan* for each sampling time-point at the focal site (KBS).

To identify specific genetic loci associated with microbiome community structure, we examined genome-wide associations (GWA) between SNPs and community structure, represented as the second axis from our NMDS analysis (described above). We did not use the first axis, since that clearly clustered with sampling date (Figure 1). We performed GWA using the *switchgrassGWAS*^*27*^ package and the same SNPs as we used in Mantel tests. To correct for population structure, we included a single variate decomposition (SVD) of pairwise genetic distance as a covariate in the linear models. The *switchgrassGWAS* package implements linear regression tests for each SNP using the *big_univLinReg* function in *bigstatsR*, which rapidly applies statistical tests across filebacked big matrices using memory mapping^98^. We calculated both a 5% false discovery rate threshold, as well as a Bonferroni-corrected p-value threshold to distinguish outlier SNPs.

To verify outlier SNPs, we examined expression-level differences of adjacent genes across divergent genotypes with RNA sequencing data from a separate study (pre-publication access through the Department of Energy Joint Genome Institute). We tested for expression differences across switchgrass genotypes using likelihood ratio tests in *DESeq2*^*99*^. We tested for expression differences across genotypes separately, and additionally examined the influence of site using a combined test for genotype ✕ site interaction.

### Important taxa

To identify OTUs that are important in structuring the fungal community, we used several complementary methods. In addition to identifying taxa important in temporal dynamics as described above, we also identified “core” taxa^15^. We examined core community taxa using custom scripts^15,100^. Core taxa are defined as those with relatively high occupancy and abundance across all samples, and represent those taxa most likely to have a close symbiosis with the host^101^. To calculate the core, we ranked OTUs by frequency, then selected all the OTUs up to the last OTU that adds a 2% increase in beta diversity (Bray-Curtis similarity) between groups ^101^. For the overall core group, we used the intersection between the core across subpopulations and the core across time. Within this core group, we used network analysis implemented in *SpiecEasi* ^102^ and *igraph*^*103*^ to build covariance networks over time. Nodes in covariance networks can be assigned to four possible groups based on the ratio of their within-module (Zi) and between-module connectivity (Pi)^104^. Those with high Zi and Pi are widely connected “network hubs”, those with low Zi and Pi are disconnected “peripherals.” Nodes with high Pi and low Zi are “connectors”, whereas those with high Zi and low Pi are “module hubs”^104^.

We used indicator species analysis to identify taxa associated with fungal rust disease. Indicator species analysis identifies particular taxa that are overrepresented based on a factor, and thus represent a useful indicator for that factor^105^. By comparing species present on infected versus uninfected leaves, we could isolate both OTUs associated with disease symptoms and those overrepresented in healthy leaves.

We further identified an *a priori* list of taxa that we expected to play important ecological roles in the phyllosphere. These included pathogens that we have previously identified in these plots, including *Puccinia* spp.^*28*^, *Bipolaris* spp., *Tilletia maclaganii*^*106*^, *and Colletotrichum spp*., and taxa with roles in herbivore prevention, including *Claviceps* spp.^*42*^, *Beauvaria* spp., and *Metarhizium* spp.^*107*^

## Data availability

Raw sequence biosamples for microbiome data have been submitted to the Sequence Read Archive^108^ as BioProject PRJNA717293. Switchgrass genetic data is available at PRJNA622568. Data analysis scripts are available on github: github.com/avanwallendael/phyllos_analysis.

## Acknowledgements

We would like to thank Lisa Vormwald, Connor Lamb, Mauricio Swartzenruber, Sydney Burtovoy, Perla Duberney, Todd Bortnem, Hannah Wilson, John Wrath, Nick Ryan, Jason Bonnette, Kate Walter, Darlene Brennan, and other field staff for their hard work at switchgrass sites. Joe Edwards, Alice MacQueen, Gary Bergstrom, Ashley Shade, and Keara Grady helped immensely with the planning for the study. Anne-Sophie Bohrer, Xingxing Li, Nate Emery, Kathy Toll, Ian Willick, Ali Soltani, and Xiaoyu Weng all helped with gathering and processing samples. We thank the Department of Energy Joint Genome Institute and collaborators, especially Kerrie Barry and Anna Lipzen, for pre-publication access to RNA-sequencing-based transcript counts and analysis of candidate genes through the gene atlas project for switchgrass. The plant material is based upon work supported in part by the Great Lakes Bioenergy Research Center, U.S. Department of Energy, Office of Science, Office of Biological and Environmental Research under Award Number DE-SC0018409. Support for this research was provided by the National Science Foundation Long-term Ecological Research Program (DEB 1832042) at the Kellogg Biological Station and by Michigan State University AgBioResearch. We received further funding from the U.S. Department of Energy through grants DE-SC0014156 to TEJ and DE-SC0017883 to DBL and National Science PGRP Award IOS1402393 to J.T.L. The work conducted by the U.S. Department of Energy Joint Genome Institute is supported by the Office of Science of the U.S. Department of Energy under Contract No. DE-AC02-05CH11231.

## Author contributions

AV designed the experiment, collected and analyzed data, and wrote the manuscript. GMNB and PBC contributed to experimental design, data analysis, and manuscript writing. LF collected data and revised the manuscript. AS, JTL, and TEJ contributed data, analyzed data, and revised the manuscript. DBL and GB contributed to experimental design and revised the manuscript.

