## Supplement for "Host genetic control of succession in the switchgrass leaf fungal microbiome"

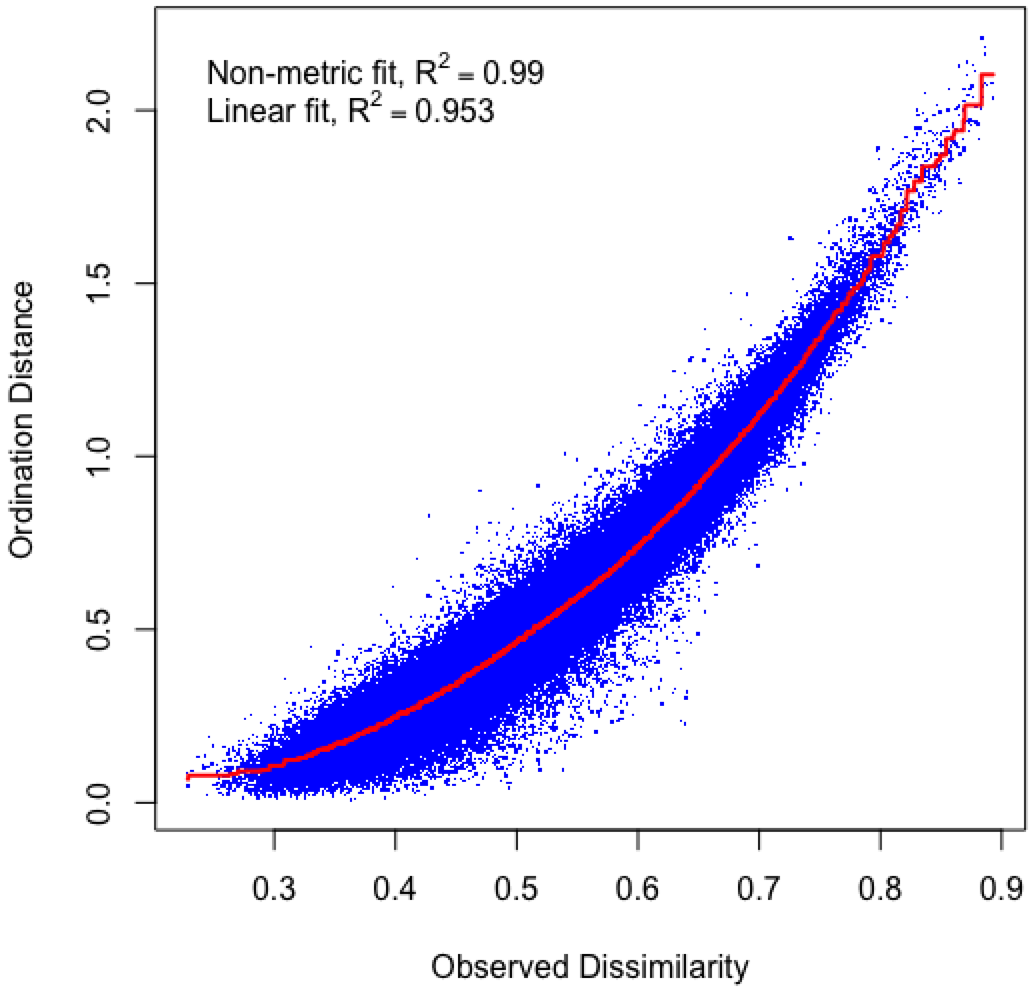


**Figure S1**: Shepard stress plot for NMDS of Kellogg Biological Station site.

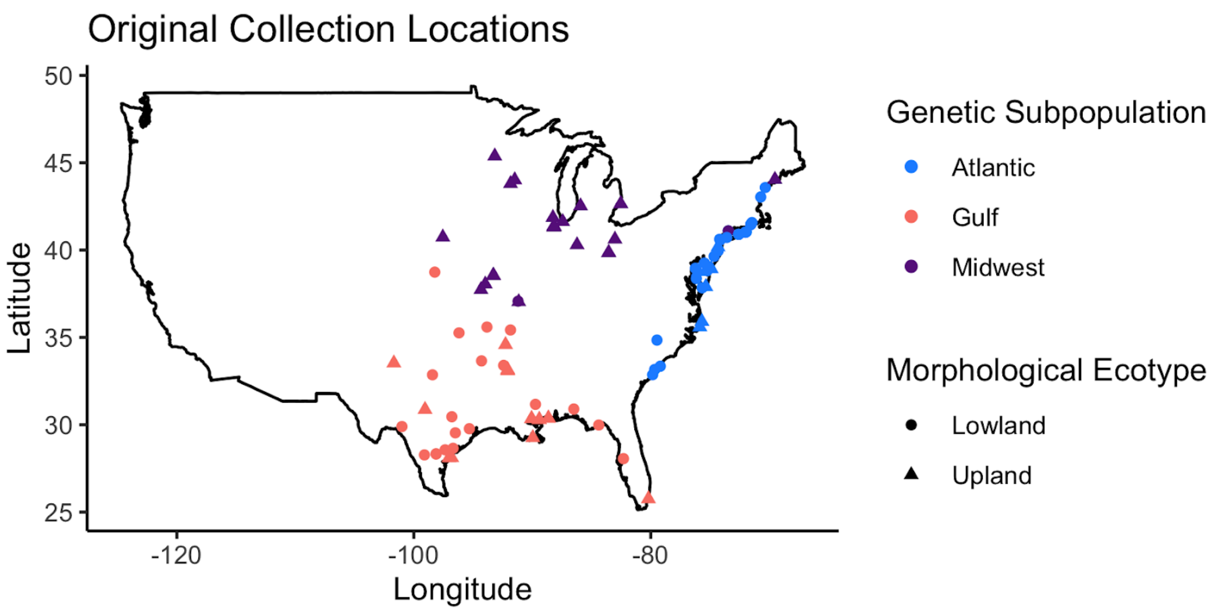


**Figure S2**: Original collection locations for samples. Latitude and Longitude are available in Table S1.

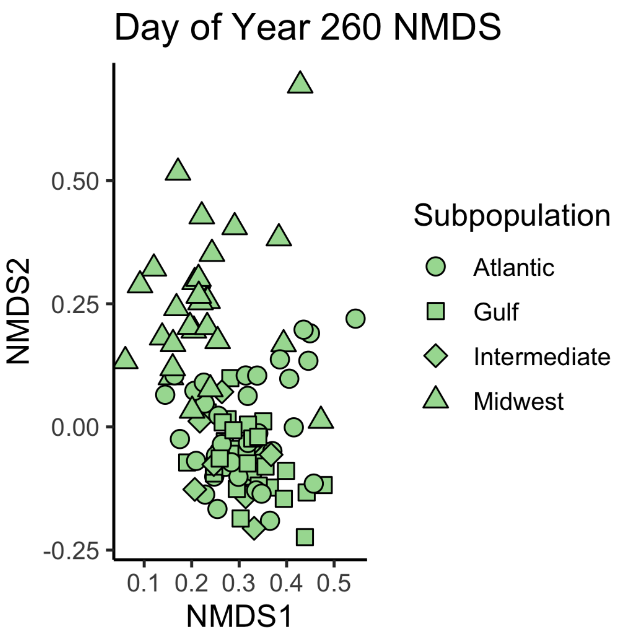


Figure S3: NMDS plot of the subsetted data used in GWAS analysis

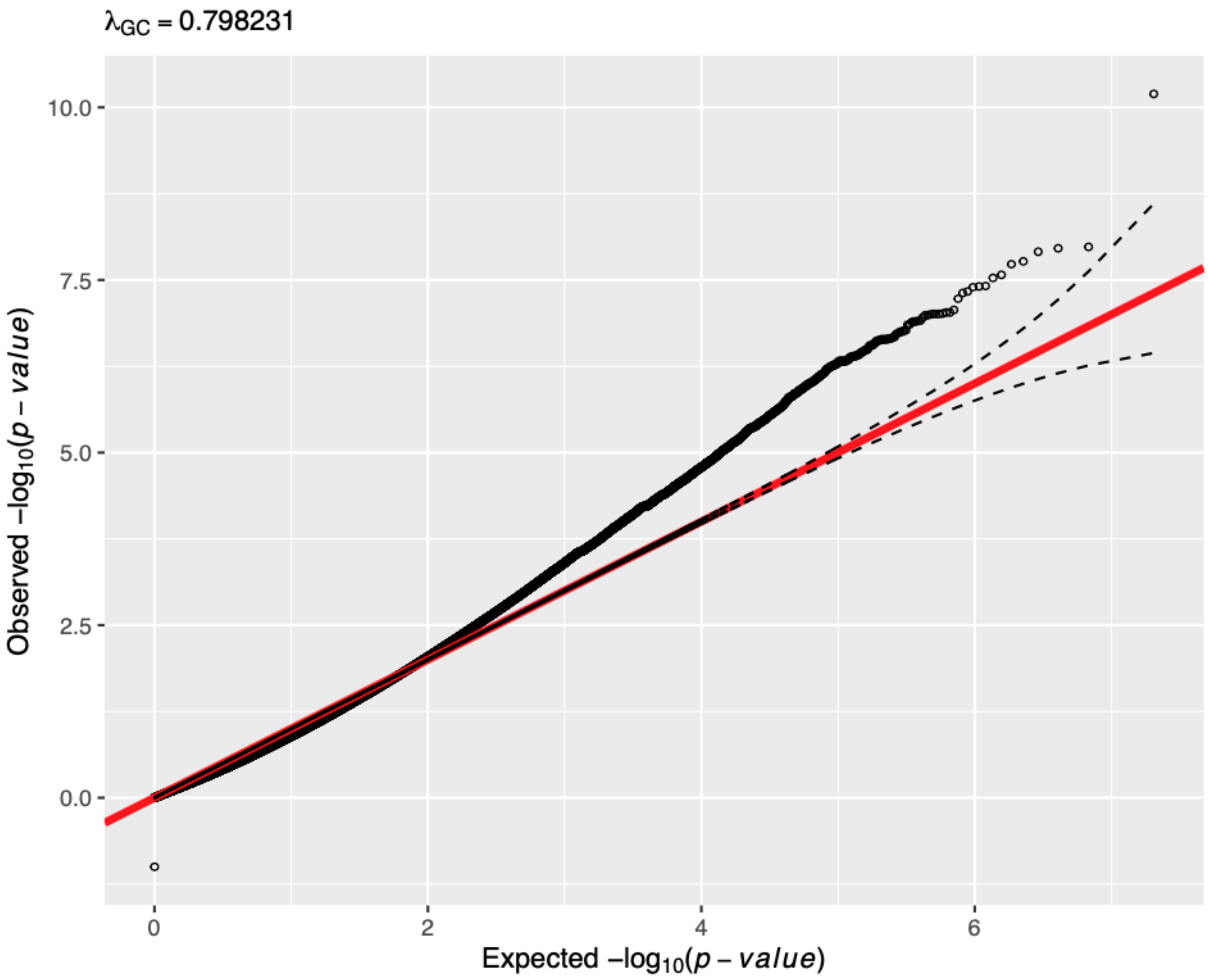


**Figure S3**: Quantile-Quantile plot for Microbiome GWAS results showing an excess of observed low p-values.

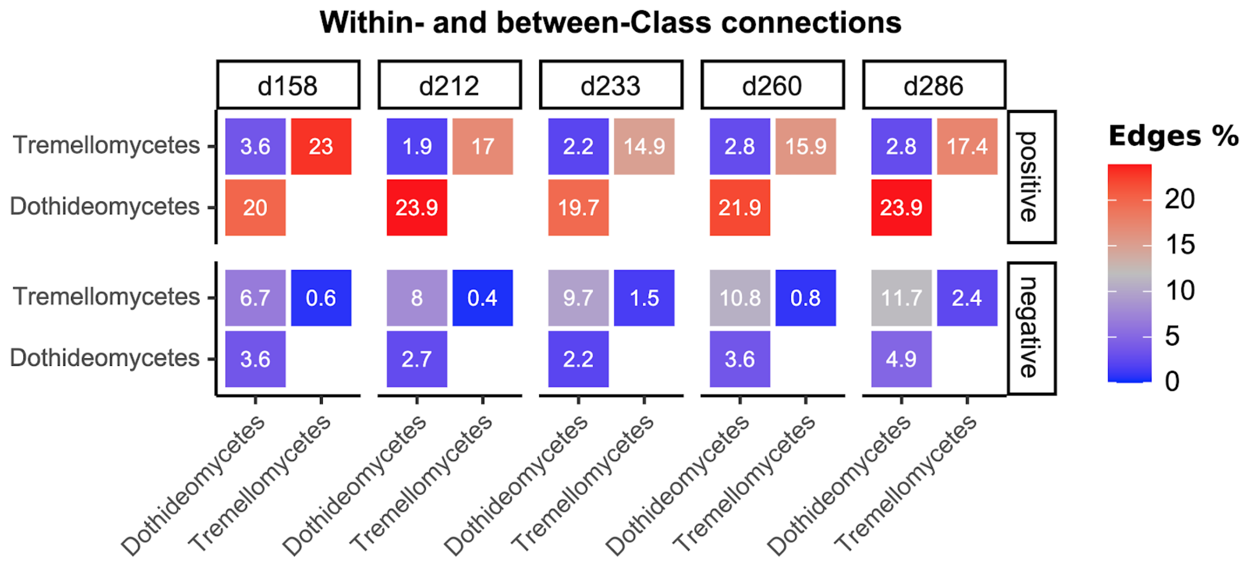


**Figure S4:** Class-level comparison of the proportion of edges linking OTUs within or between each Class in a time point. Column facets show day of year (DOY), and row facets show whether connections were negative or positive.

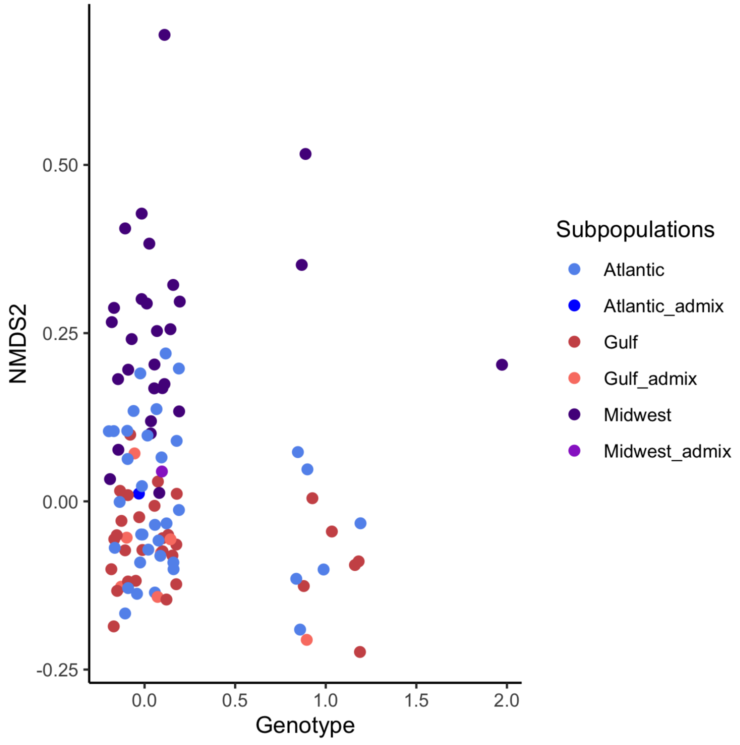


**Figure S5:**Phenotypic (NMDS2) values for the outlier SNP, Chr02N_57831909. The x-axis shows jittered genotypic value, with 0 and 2 as homozygotes, and 1 as the heterozygote. Points are colored by population. Subpopulations with the _admix suffix show substantial admixture from other populations.

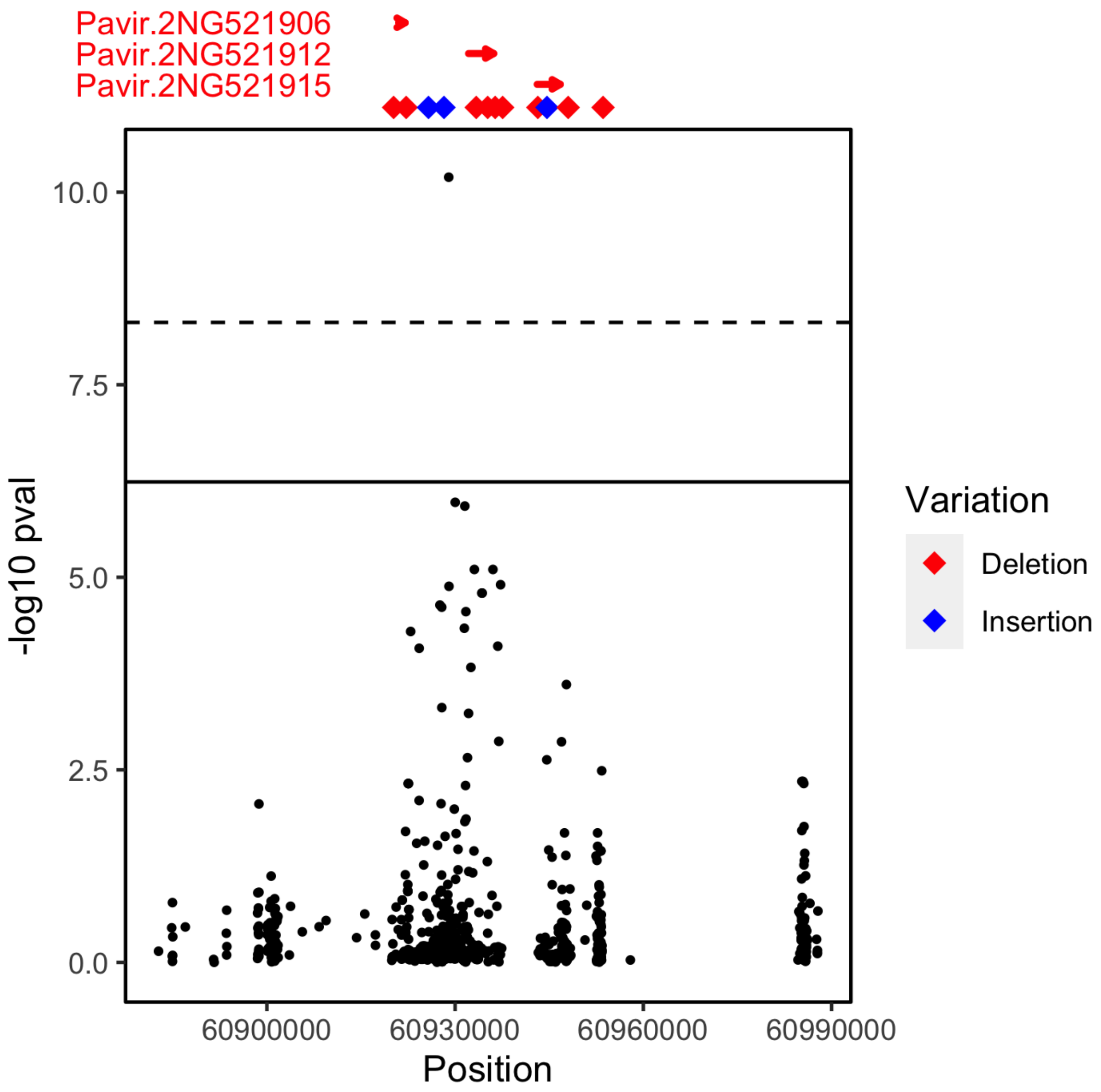


**Figure S6:** Insertions and deletions in the region of our outlier SNP were overrepresented in the area surrounding the candidate genes, indicating structural variation across genotypes that may account for genetic and phenotypic differences. Red diamonds show deletions, and blue triangles show insertions. Red arrows with text show nearby genes.

**Table S1**: Core microbiome taxonomy and functional guilds

| OTU_ID | Phylum | Class | Genus | Species | BLAST Percent ID | Functional Guild | Guild Source |
|---|---|---|---|---|---|---|---|
| OTU_2 | Ascomycota | Dothideomycetes | Alternaria | NA | 100 | Pathogen | Source |

| | | | | | | | |
|---|---|---|---|---|---|---|---|
| OTU_32 | Ascomycota | Dothideomycetes | Alternaria | NA | 100 | Pathogen | Source |
| OTU_6104 | Ascomycota | Dothideomycetes | Aureobasidium | NA | 96.094 | Yeast or Yeast-like | Source |
| OTU_1760 | Basidiomycota | Tremellomycetes | Bulleromyces | Bulleromyces albus | 0 | Yeast or Yeast-like | Source |
| OTU_5936 | Basidiomycota | Tremellomycetes | Bulleromyces | Bulleromyces albus | 0 | Yeast or Yeast-like | Source |
| OTU_738 | Basidiomycota | Tremellomycetes | Bulleromyces | Bulleromyces albus | 99.422 | Yeast or Yeast-like | Source |
| OTU_392 | Basidiomycota | Tremellomycetes | Bulleromyces | Bulleromyces albus | 0 | Yeast or Yeast-like | Source |
| OTU_47 | Ascomycota | Dothideomycetes | Cladosporium | Cladosporium grevilleae | 100 | saprobe | Source |
| OTU_21 | Ascomycota | Dothideomycetes | Coniothyrium | Coniothyrium | 100 | Other Pathogen | Source |
| OTU_380 | Basidiomycota | Tremellomycetes | Cryptococcus | NA | 100 | Yeast or Yeast-like | Source |
| OTU_2137 | Ascomycota | Dothideomycetes | Didymella | NA | 99.471 | Other Pathogen | Source |
| OTU_877 | Basidiomycota | Tremellomycetes | Dioszegia | Dioszegia athyri | 98.919 | Yeast or Yeast-like | Source |
| OTU_1303 | Basidiomycota | Tremellomycetes | Dioszegia | Dioszegia athyri | 98.413 | Yeast or Yeast-like | Source |
| OTU_1615 | Basidiomycota | Tremellomycetes | Dioszegia | Dioszegia athyri | 98.919 | Yeast or Yeast-like | Source |
| OTU_464 | Basidiomycota | Tremellomycetes | Dioszegia | Dioszegia hungarica | 0 | Yeast or Yeast-like | Source |
| OTU_1011 | Basidiomycota | Tremellomycetes | Dioszegia | Dioszegia hungarica | 98.817 | Yeast or Yeast-like | Source |
| OTU_5498 | Basidiomycota | Tremellomycetes | Dioszegia | Dioszegia hungarica | 0 | Yeast or Yeast-like | Source |
| OTU_822 | Basidiomycota | Tremellomycetes | Dioszegia | NA | 100 | Yeast or Yeast-like | Source |
| OTU_1367 | Basidiomycota | Tremellomycetes | Dioszegia | NA | 100 | Yeast or Yeast-like | Source |
| OTU_414 | Basidiomycota | Tremellomycetes | Dioszegia | NA | 100 | Yeast or Yeast-like | Source |
| OTU_2213 | Basidiomycota | Tremellomycetes | Dioszegia | NA | 100 | Yeast or Yeast-like | Source |
| OTU_677 | Ascomycota | Dothideomycetes | Dissoconium | Dissoconium eucalypti | 99 | Other Pathogen | Source |
| OTU_18 | Ascomycota | Dothideomycetes | Dissoconium | NA | 98.02 | Other Pathogen | |
| OTU_35 | Ascomycota | Dothideomycetes | Epicoccum | Epicoccum dendrobii | 100 | Mycoparasite | Source |
| OTU_165 | Ascomycota | Dothideomycetes | Epicoccum | Epicoccum dendrobii | 97.98 | Mycoparasite | Source |
| OTU_2231 | Basidiomycota | Cystobasidiomycetes | Erythrobasidium | Erythrobasidium hasegawianum | 97.959 | Yeast or Yeast-like | Source |
| OTU_770 | Basidiomycota | Cystobasidiomycetes | Erythrobasidium | NA | 98.469 | Yeast or Yeast-like | Source |
| OTU_52 | Basidiomycota | Cystobasidiomycetes | Erythrobasidium | Erythrobasidium yunnanense | 98.985 | Yeast or Yeast-like | Source |
| OTU_33 | Basidiomycota | Tremellomycetes | Filobasidium | Filobasidium floriforme | 99 | Yeast or Yeast-like | Source |
| OTU_980 | Basidiomycota | Tremellomycetes | Filobasidium | NA | 0 | Yeast or Yeast-like | Source |
| OTU_1226 | Basidiomycota | Tremellomycetes | Filobasidium | Filobasidium wieringae | 100 | Yeast or Yeast-like | Source |

| | | | | | | | |
|---|---|---|---|---|---|---|---|
| OTU_37 | Ascomycota | Sordariomycetes | Fusarium | Fusarium sporotrichioides | 100 | Pathogen | Source |
| OTU_2922 | Basidiomycota | Tremellomycetes | Hannaella | Hannaella sinensis | 0 | Yeast or Yeast-like | Source |
| OTU_791 | Basidiomycota | Tremellomycetes | Hannaella | Hannaella sinensis | 0 | Yeast or Yeast-like | Source |
| OTU_1724 | Basidiomycota | Tremellomycetes | Hannaella | Hannaella sinensis | 0 | Yeast or Yeast-like | Source |
| OTU_14 | Ascomycota | Dothideomycetes | Keissleriella | Keissleriella caraganae | 100 | Other Pathogen | Source |
| OTU_92 | Basidiomycota | Agaricostilbomycetes | Kondoa | Kondoa | 97.5 | Yeast or Yeast-like | Source |
| OTU_24 | Basidiomycota | Agaricostilbomycetes | Kondoa | Kondoa | 99 | Yeast or Yeast-like | Source |
| OTU_38 | Basidiomycota | Agaricostilbomycetes | Kondoa | Kondoa miscanthi | 99.502 | Yeast or Yeast-like | Source |
| OTU_28 | Basidiomycota | Agaricostilbomycetes | Kondoa | NA | 94.634 | Yeast or Yeast-like | Source |
| OTU_72 | Ascomycota | Dothideomycetes | Leptospora | Leptospora | 99 | | |
| OTU_6 | Ascomycota | Sordariomycetes | Microdochium | Microdochium seminicola | 95.05 | Pathogen | Source |
| OTU_158 | Ascomycota | Dothideomycetes | Mycosphaerella | Mycosphaerella tassiana | 99.5 | Pathogen | Source |
| OTU_7 | Ascomycota | NA | NA | NA | 0 | | |
| OTU_426 | Ascomycota | Dothideomycetes | NA | NA | 98.942 | | |
| OTU_1076 | Ascomycota | Dothideomycetes | NA | NA | 99.474 | | |
| OTU_1690 | Ascomycota | Dothideomycetes | NA | NA | 98.454 | | |
| OTU_42 | Ascomycota | Dothideomycetes | NA | NA | 0 | | |
| OTU_5 | Ascomycota | Dothideomycetes | NA | NA | 100 | | |
| OTU_103 | NA | NA | NA | NA | 100 | | |
| OTU_115 | NA | NA | NA | NA | 99.5 | | |
| OTU_15 | Ascomycota | Leotiomycetes | NA | NA | 89.447 | | |
| OTU_93 | Basidiomycota | Microbotryomycetes | NA | NA | 0 | | |
| OTU_56 | Ascomycota | Sordariomycetes | NA | NA | 100 | | |
| OTU_341 | Ascomycota | Dothideomycetes | NA | NA | 98.947 | | |
| OTU_1207 | Ascomycota | Dothideomycetes | NA | NA | 98.469 | | |
| OTU_1723 | Ascomycota | Dothideomycetes | NA | NA | 99.479 | | |
| OTU_610 | Ascomycota | Dothideomycetes | NA | NA | 99.471 | | |
| OTU_20 | Ascomycota | Dothideomycetes | NA | NA | 100 | | |
| OTU_79 | Ascomycota | Dothideomycetes | NA | NA | 99.5 | | |
| OTU_705 | Ascomycota | Dothideomycetes | NA | NA | 98.953 | | |
| OTU_2655 | Ascomycota | Dothideomycetes | NA | NA | 100 | | |
| OTU_1041 | Ascomycota | Dothideomycetes | NA | NA | 97 | | |
| OTU_11 | Ascomycota | Dothideomycetes | NA | NA | 100 | | |
| OTU_8 | Ascomycota | Dothideomycetes | NA | NA | 100 | | |

| | | | | | | | |
|---|---|---|---|---|---|---|---|
| OTU_13 | Ascomycota | Dothideomycetes | NA | NA | 100 | | |
| OTU_5444 | Ascomycota | Dothideomycetes | NA | NA | 97.015 | | |
| OTU_23 | Ascomycota | Leotiomycetes | NA | NA | 100 | | |
| OTU_628 | Ascomycota | Leotiomycetes | NA | NA | 96.5 | | |
| OTU_248 | Basidiomycota | Tremellomycetes | NA | NA | 94.767 | | |
| OTU_505 | Basidiomycota | Tremellomycetes | NA | NA | 94.35 | | |
| OTU_1234 | Basidiomycota | Tremellomycetes | NA | NA | 94.767 | | |
| OTU_3488 | Basidiomycota | Tremellomycetes | NA | NA | 93.605 | | |
| OTU_4633 | Basidiomycota | Tremellomycetes | NA | NA | 94.186 | | |
| OTU_6081 | Basidiomycota | Tremellomycetes | NA | NA | 94.767 | | |
| OTU_5180 | Basidiomycota | Tremellomycetes | NA | NA | 94.767 | | |
| OTU_41 | Basidiomycota | Tremellomycetes | NA | NA | 95.567 | | |
| OTU_3292 | Ascomycota | Dothideomycetes | Neoascochyta | Neoascochyta exitialis | 99.487 | Pathogen | Source |
| OTU_443 | Ascomycota | Dothideomycetes | Neoascochyta | NA | 98.953 | Pathogen | Source |
| OTU_429 | Ascomycota | Dothideomycetes | Neoascochyta | NA | 100 | Pathogen | Source |
| OTU_340 | Ascomycota | Dothideomycetes | Neodevriesia | Neodevriesia poagena | 100 | | Source |
| OTU_61 | Ascomycota | Sordariomycetes | Nigrospora | Nigrospora oryzae | 100 | Pathogen | Source |
| OTU_669 | Basidiomycota | Tremellomycetes | Papiliotrema | Papiliotrema pseudoalba | 0 | Yeast or Yeast-like | Source |
| OTU_6048 | Basidiomycota | Tremellomycetes | Papiliotrema | Papiliotrema pseudoalba | 0 | Yeast or Yeast-like | Source |
| OTU_1993 | Basidiomycota | Tremellomycetes | Papiliotrema | NA | 100 | Yeast or Yeast-like | Source |
| OTU_394 | Basidiomycota | Tremellomycetes | Papiliotrema | NA | 100 | Yeast or Yeast-like | Source |
| OTU_7324 | Basidiomycota | Tremellomycetes | Papiliotrema | NA | 99.425 | Yeast or Yeast-like | Source |
| OTU_6867 | Basidiomycota | Tremellomycetes | Papiliotrema | NA | 100 | Yeast or Yeast-like | Source |
| OTU_4109 | Basidiomycota | Tremellomycetes | Papiliotrema | NA | 98.851 | Yeast or Yeast-like | Source |
| OTU_5837 | Basidiomycota | Tremellomycetes | Papiliotrema | NA | 98.851 | Yeast or Yeast-like | Source |
| OTU_4177 | Basidiomycota | Tremellomycetes | Papiliotrema | NA | 98.857 | Yeast or Yeast-like | Source |
| OTU_74 | Ascomycota | Dothideomycetes | Paraophiobolus | Paraophiobolus arundinis | 100 | | Source |
| OTU_49 | Ascomycota | Dothideomycetes | Paraphaeosphaeria | Paraphaeosphaeria michotii | 100 | | Source |
| OTU_137 | Basidiomycota | Agaricomycetes | Peniophora | NA | 99 | | Source |
| OTU_43 | Ascomycota | Dothideomycetes | Phaeosphaeria | Phaeosphaeria | 99.497 | Pathogen | Source |
| OTU_125 | Ascomycota | Dothideomycetes | Phaeosphaeria | Phaeosphaeria | 100 | Pathogen | |

| | | | | | | | |
|---|---|---|---|---|---|---|---|
| OTU_83 | Ascomycota | Dothideomycetes | Phaeosphaeria | NA | 92.04 | Pathogen | |
| OTU_232 | Ascomycota | Dothideomycetes | Phaeosphaeria | NA | 95.855 | | |
| OTU_3 | Ascomycota | Dothideomycetes | Phoma | NA | 100 | | Source |
| OTU_59 | Ascomycota | Dothideomycetes | Ramularia | NA | 0 | | Source |
| OTU_1684 | Basidiomycota | Tremellomycetes | Saitozyma | Saitozyma paraflava | 0 | | Source |
| OTU_1028 | Basidiomycota | Tremellomycetes | Saitozyma | Saitozyma paraflava | 0 | | Source |
| OTU_50 | Ascomycota | Sordariomycetes | Sarocladium | NA | 100 | | Source |
| OTU_91 | Ascomycota | Dothideomycetes | Septoria | NA | 100 | | Source |
| OTU_143 | Ascomycota | Dothideomycetes | Setomelanomma | Setomelanomma | 0 | | Source |
| OTU_4 | Ascomycota | Dothideomycetes | Sphaerellopsis | Sphaerellopsis filum | 97 | Mycoparasite | Source |
| OTU_31 | Basidiomycota | Microbotryomycetes | Sporobolomyces | Sporobolomyces patagonicus | 98.985 | Yeast or Yeast-like | Source |
| OTU_17 | Basidiomycota | Microbotryomycetes | Sporobolomyces | Sporobolomyces phaffii | 100 | Yeast or Yeast-like | Source |
| OTU_12 | Basidiomycota | Microbotryomycetes | Sporobolomyces | Sporobolomyces roseus | 99.5 | | Source |
| OTU_30 | Ascomycota | Dothideomycetes | Stagonospora | Stagonospora pseudovitensis | 98 | | Source |
| OTU_71 | Basidiomycota | Cystobasidiomycetes | Symmetrospora | Symmetrospora coprosmae | 100 | | Source |
| OTU_476 | Basidiomycota | Cystobasidiomycetes | Symmetrospora | Symmetrospora gracilis | 100 | | Source |
| OTU_86 | Basidiomycota | Cystobasidiomycetes | Symmetrospora | NA | 99.479 | | Source |
| OTU_1663 | Basidiomycota | Cystobasidiomycetes | Symmetrospora | NA | 99.49 | | Source |
| OTU_22 | Ascomycota | Taphrinomycetes | Taphrina | Taphrina | 100 | Pathogen | Source |
| OTU_64 | Ascomycota | Taphrinomycetes | Taphrina | Taphrina communis | 100 | Pathogen | Source |
| OTU_16 | Ascomycota | Taphrinomycetes | Taphrina | Taphrina confusa | 99 | Pathogen | Source |
| OTU_26 | Ascomycota | Taphrinomycetes | Taphrina | Taphrina letifera | 99.5 | Pathogen | Source |
| OTU_10 | Ascomycota | Taphrinomycetes | Taphrina | NA | 87.624 | Pathogen | Source |
| OTU_54 | Ascomycota | Taphrinomycetes | Taphrina | Taphrina tormentillae | 99.5 | Pathogen | Source |
| OTU_105 | Basidiomycota | Exobasidiomycetes | Tilletiopsis | Tilletiopsis washingtonensis | 98 | | Source |
| OTU_75 | Basidiomycota | Tremellomycetes | Udeniomyces | Udeniomyces pyricola | 100 | | Source |
| OTU_9 | Ascomycota | Dothideomycetes | Uwebraunia | Uwebraunia dekkeri | 100 | | Source |
| OTU_4086 | Ascomycota | Dothideomycetes | Vagicola | Vagicola chlamydospora | 96.535 | | Source |
| OTU_2473 | Basidiomycota | Tremellomycetes | Vishniacozyma | Vishniacozyma victoriae | 96.196 | | Source |
| OTU_1571 | Basidiomycota | Tremellomycetes | Vishniacozyma | Vishniacozyma victoriae | 98.37 | | Source |
| OTU_3311 | Basidiomycota | Tremellomycetes | Vishniacozyma | Vishniacozyma victoriae | 100 | | Source |
| OTU_979 | Ascomycota | Dothideomycetes | Zymoseptoria | NA | 96.891 | | Source |

**Table S2**: Sample information. Ecotype and subpopulation membership are estimated using SNP data. Latitude and longitude denote original collection site, if known.

| PLOT_ID | PLANT_ID | Ecotype_SNP | Subpopulation | Latitude | Longitude |
|---|---|---|---|---|---|
| M1001 | J208.B | Lowland | eastcoast_admixed | 34.845 | -79.470 |
| M1005 | J297.A | Lowland | Texas | NA | NA |
| M1010 | J427.B | Lowland | eastcoast | 40.620 | -74.180 |
| M1107 | J585.A | Lowland | eastcoast | 32.865 | -79.837 |
| M1116 | J294.A | Upland | Texas_admixed | 28.110 | -97.029 |
| M1120 | J368.B | Upland | midwest | 42.528 | -85.922 |
| M1202 | J344.C | Upland | midwest | 41.305 | -88.172 |
| M1209 | J330.A | Lowland | Texas | 28.276 | -99.101 |
| M1218 | J499.A | Upland | Gulfcoast | 30.348 | -90.057 |
| M1401 | J022.B | Lowland | Texas | 28.333 | -98.118 |
| M1407 | J186.A | Lowland | Gulfcoast_admixed | 29.986 | -84.387 |
| M1408 | J491.B | Upland | midwest | 38.050 | -93.967 |
| M1411 | J661.A | Lowland | eastcoast | 37.841 | -75.654 |
| M1417 | J589.B | Upland | eastcoast | 37.907 | -75.351 |
| M1420 | J314.A | Lowland | Texas_admixed | 29.775 | -95.310 |
| M1602 | J483.C | Upland | Gulfcoast | 33.085 | -92.070 |
| M1605 | J448.A | Upland | midwest | NA | NA |
| M1801 | J028.C | Lowland | Texas | 35.593 | -93.825 |
| M1804 | J461.C | Upland | Gulfcoast | 30.299 | -89.407 |
| M1806 | J496.A | Upland | Gulfcoast | 29.262 | -89.952 |
| M2002 | J462.C | NA | NA | 30.481 | -92.668 |
| M2005 | J306.A | Upland | Texas | 30.874 | -99.050 |
| M2015 | J433.B | Lowland | eastcoast | 41.020 | -72.010 |
| M2018 | J614.B | Upland | eastcoast | 38.935 | -74.906 |
| M2020 | J482.B | Upland | Gulfcoast | 33.150 | -92.076 |
| M2108 | J502.C | Upland | Gulfcoast | 30.377 | -88.634 |
| M2201 | J538.C | Lowland | eastcoast | 41.017 | -72.001 |
| M2214 | J305.A | Upland | Texas_admixed | 28.117 | -96.800 |
| M2216 | J396.A | Upland | midwest | 43.800 | -91.830 |
| M2217 | J355.A | Upland | midwest | 41.347 | -88.139 |
| M2218 | J022.C | Lowland | Texas | 28.333 | -98.118 |
| M2401 | J672.A | Lowland | eastcoast | 39.255 | -75.466 |
| M2402 | J598.B | Upland | eastcoast | 38.772 | -75.977 |
| M2417 | J348.C | Upland | midwest | 40.620 | -83.021 |
| M2601 | J481.A | Lowland | Gulfcoast_admixed | 33.397 | -92.414 |
| M2607 | J536.C | Lowland | eastcoast | 41.104 | -73.451 |
| M2614 | J222.A | Lowland | Texas | 30.460 | -96.785 |
| M2620 | J682.C | Upland | eastcoast | 40.179 | -74.317 |
| M2802 | J315.A | Lowland | Texas | NA | NA |
| M2806 | BLK.17 | NA | NA | NA | NA |
| M2920 | J331.A | Lowland | Texas | 29.538 | -96.493 |
| M3001 | J299.A | Lowland | Texas_admixed | NA | NA |

| | | | | | |
|---|---|---|---|---|---|
| M3003 | J663.B | Lowland | eastcoast | 38.374 | -76.149 |
| M3007 | BLK.20 | NA | NA | NA | NA |
| M3012 | J502.C | Upland | Gulfcoast | 30.377 | -88.634 |
| M3201 | J308.A | Lowland | Texas_admixed | 28.567 | -97.352 |
| M3202 | J534.C | Lowland | eastcoast | 40.586 | -74.119 |
| M3205 | J580.A | Upland | midwest | 41.861 | -88.254 |
| M3209 | BLK.21 | NA | NA | NA | NA |
| M3210 | J657.C | Lowland | midwest | 37.073 | -91.197 |
| M3220 | J582.C | Upland | midwest | 41.365 | -88.187 |
| M3520 | J461.B | Upland | Gulfcoast | 30.299 | -89.407 |
| M3606 | J245.A | Lowland | Texas | 32.860 | -98.420 |
| M3613 | J065.A | Lowland | Texas | 28.060 | -82.300 |
| M3615 | J651.C | Lowland | Texas | NA | NA |
| M3618 | J635.A | Lowland | eastcoast | 43.580 | -70.329 |
| M3803 | J353.B | Upland | midwest | 39.854 | -83.531 |
| M3819 | J318.A | Lowland | Texas | 28.654 | -96.681 |
| M4001 | J484.B | Lowland | Texas | 33.658 | -94.281 |
| M4017 | J379.B | Upland | midwest | 42.648 | -82.529 |
| M4021 | J673.D | Lowland | eastcoast | 38.820 | -75.228 |
| M4209 | J390.B | Upland | midwest | 40.300 | -86.220 |
| M4210 | J016.C | Lowland | Texas | 35.260 | -96.180 |
| M4211 | J018.C | Lowland | Texas | 35.427 | -91.837 |
| M4410 | J504.C | Lowland | Gulfcoast | 31.170 | -89.730 |
| M4412 | J024.A | Upland | midwest | 51.219 | 4.402 |
| M4416 | J004.B | Lowland | Texas | 38.737 | -98.228 |
| M4418 | J355.C | Upland | midwest | 41.347 | -88.139 |
| M4419 | J416.B | Lowland | eastcoast | 40.600 | -74.130 |
| M4520 | J499.C | Upland | Gulfcoast | 30.348 | -90.057 |
| M4602 | J656.B | Upland | midwest | 37.044 | -91.141 |
| M4610 | J536.A | Lowland | midwest_admixed | 41.104 | -73.451 |
| M4617 | J482.A | Upland | Gulfcoast | 33.150 | -92.076 |
| M4802 | J660.C | Lowland | Texas | 29.898 | -100.997 |
| M4805 | J538.A | Lowland | eastcoast | 41.017 | -72.001 |
| M4817 | J614.A | Upland | eastcoast | 38.935 | -74.906 |
| M5401 | J341.A | Upland | Texas | 33.533 | -101.680 |
| M5404 | J352.B | Upland | midwest | 39.839 | -83.573 |
| M5405 | J045.C | Upland | midwest | 40.740 | -97.546 |
| M5409 | J650.A | Upland | Texas_admixed | 25.762 | -80.192 |
| M5413 | J646.C | Lowland | eastcoast | 41.463 | -71.554 |
| M5418 | J489.A | Upland | midwest | 37.737 | -94.328 |
| M5804 | J419.A | Lowland | Texas | NA | NA |
| M5809 | J387.C | Upland | midwest | 38.551 | -93.299 |
| M5810 | J477.C | Upland | Gulfcoast_admixed | 34.587 | -92.254 |
| M5811 | BLK.50 | NA | NA | NA | NA |
| M5812 | J386.C | Upland | midwest | 38.548 | -93.258 |
| M5813 | J596.C | Lowland | eastcoast | 38.952 | -76.234 |

| | | | | | |
|---|---|---|---|---|---|
| M5817 | J258.A | Upland | midwest | 45.382 | -93.164 |
| M6005 | J636.C | Upland | midwest | 44.043 | -69.516 |
| M6012 | J593.B | Upland | eastcoast | 35.600 | -75.848 |
| M6016 | J522.B | Lowland | eastcoast | 39.644 | -74.644 |
| M6019 | J613.C | Upland | eastcoast | 35.908 | -75.676 |
| M6204 | J521.A | Lowland | eastcoast | 39.965 | -74.312 |
| M6215 | J602.C | Lowland | eastcoast | 33.352 | -79.194 |
| M6221 | J344.A | Upland | midwest | 41.305 | -88.172 |
| M6607 | J646.A | Lowland | eastcoast | 41.463 | -71.554 |
| M6608 | J607.A | Lowland | eastcoast | 41.575 | -71.454 |
| M6610 | J521.C | Lowland | eastcoast | 39.965 | -74.312 |
| M6611 | J609.C | Lowland | eastcoast | 33.167 | -79.666 |
| M6613 | J432.B | Lowland | eastcoast | 41.040 | -71.930 |
| M6614 | J385.A | Upland | midwest | 38.532 | -93.291 |
| M6615 | J634.A | Lowland | eastcoast | 43.038 | -70.716 |
| M6705 | J646.B | Lowland | eastcoast | 41.463 | -71.554 |
| M6706 | J428.B | Lowland | eastcoast | 40.720 | -73.580 |
| M6708 | J521.B | Lowland | eastcoast | 39.965 | -74.312 |
| M6711 | J403.B | Upland | midwest | 44.020 | -91.480 |
| M6712 | J525.C | Lowland | eastcoast | 40.903 | -72.571 |
| M6713 | J525.A | Lowland | eastcoast | 40.903 | -72.571 |
| M6714 | J456.A | Lowland | Gulfcoast | 30.900 | -86.494 |
| M6802 | J525.B | Lowland | eastcoast | 40.903 | -72.571 |
| M6803 | J379.A | Upland | midwest | 42.648 | -82.529 |
| M6813 | J393.B | Upland | midwest | 41.630 | -87.430 |
| M6817 | J538.B | Lowland | eastcoast | 41.017 | -72.001 |

**Table S3:** PCR conditions for ITS amplification

| Round 1 (10X) | | Round 2 (10X) | | Round 3 (15X) | |
|---|---|---|---|---|---|
| **Temp (°C)** | **Time** | **Temp (°C)** | **Time** | **Temp (°C)** | **Time** |
| 94° | 5:00 | 94° | 5:00 | 94° | 5:00 |
| 94° | 0:30 | 94° | 0:30 | 94° | 0:30 |
| **50°** | 0:30 | **50°** | 0:35 | **63°** | 0:35 |
| 72° | 1:20 | 72° | 1:20 | 72° | 1:20 |
| 72° | 7:00 | 72° | 7:00 | 72° | 7:00 |
| 4° | inf | 4° | inf | 4° | inf |

| **round 1----- 1x plate** |  |  |  |
|---|---|---|---|
| **Reagents** | **µL** | **Reactions** | **Total (µL)** |
| Dream Taq GREEN | 10.4 | 104 | 1081.6 |
| ITS1F | 0.625 | 104 | 65 |
| ITS2 | 0.625 | 104 | 65 |
| BSA 3% | 3.35 | 104 | 348.4 |
| H2O | 1 | 104 | 104 |
| **Total (µL)** | **16** | | **1,664** |
| DNA (µL) | **4** | | |
| Final reaction volume | 20 | | |
| **round 2-----1x plate** |  |  |  |
| Dream Taq GREEN | **µL** | **Reactions** | **Total (µL)** |
| master MIX | 6.25 | 104 | 650 |
| ITS1F* | 0.375 | 104 | 39 |
| ITS2* | 0.375 | 104 | 39 |
| BSA 3% | 2 | 104 | 208 |
| H2O | 2 | 104 | 208 |
| **Total (µL)** | **11** | | **1,144** |
| DNA (µL) | **2** | | |
| Final reaction volume | 13 | | |
| **round 3 ----- 1x plate** |  |  |  |
| Dream Taq GREEN | **µL** | **Reactions** | **Total (µL)** |
| master MIX | 8 | 104 | 832 |
| Forward primer F | 0.5 | 104 | 52 |
| Barcode primers R | - | - | - |
| BSA 3% | 1.5 | 104 | 156 |
| H2O | 0 | 104 | 0 |
| **Total (µL)** | **10+1ulBar** | | **1,040** |
| DNA (µL) | **4** | | |
